# Indication-specific tumor evolution and its impact on neoantigen targeting and biomarkers for individualized cancer immunotherapies

**DOI:** 10.1101/2021.03.15.434617

**Authors:** Amy A. Lo, Andrew Wallace, Daniel Oreper, Nicolas Lounsbury, Charles Havnar, Ximo Pechuan-Jorge, Thomas D. Wu, Richard Bourgon, Ryan Jones, Katrina Krogh, Guang-Yu Yang, Oliver A. Zill

**Affiliations:** Department of Research Pathology, Genentech, Inc; Department of Oncology Bioinformatics, Genentech, Inc; Department of Cancer Immunology, Genentech, Inc; Department of Pathology, Feinberg School of Medicine, Northwestern University

## Abstract

Individualized neoantigen specific immunotherapy (iNeST) requires robustly expressed clonal neoantigens for efficacy, but tumor mutational heterogeneity, loss of neoantigen expression, and variable tissue sampling present challenges. To characterize these potential obstacles, we combined multi-region sequencing (MR-seq) analysis of five untreated, synchronously sampled metastatic solid tumors with re-analysis of published MR-seq data from 103 patients. Branching evolution in colorectal cancer and renal cell carcinoma led to fewer clonal neoantigens and to clade-specific neoantigens (those shared across a subset of tumor regions but not fully clonal), with the latter not being readily distinguishable in single tumor samples. Prioritizing mutations with higher purity- and ploidy-adjusted variant allele frequency enriched for globally clonal neoantigens (those found in all tumor regions), whereas estimated cancer cell fraction derived from clustering-based tools, surprisingly, did not. Neoantigen quality was associated with loss of neoantigen expression in the bladder cancer case, and HLA-allele loss was observed in the renal and non-small cell lung cancer cases. Our results show that indication type, multi-lesion sampling, neoantigen expression, and HLA allele retention are important factors for iNeST targeting and patient selection.

## Introduction

Tumor neoantigens are mutant peptide sequences that arise from expressed somatic mutations and can mediate anti-tumor T cell responses when presented on MHC molecules. The vast majority of neoantigens correspond to private passenger mutations ^1, 2^, meaning they are unique to a given tumor and have unknown functional consequences. Several types of individualized cancer immunotherapies have been under development in recent years, with a major modality being neoantigen vaccines ^3–5^. These immunotherapies typically utilize a single tumor biopsy for identification of tumor mutations, which are then translated into mutant peptide sequences that are fed into MHC binding/presentation prediction algorithms. These predictions allow the selection of specific patient-derived tumor neoantigens, which then inform the design of a customized therapeutic (Fig. 1A) such as genetically modified neoantigen-specific T-cells or a neoantigen vaccine.

**Fig. 1:**
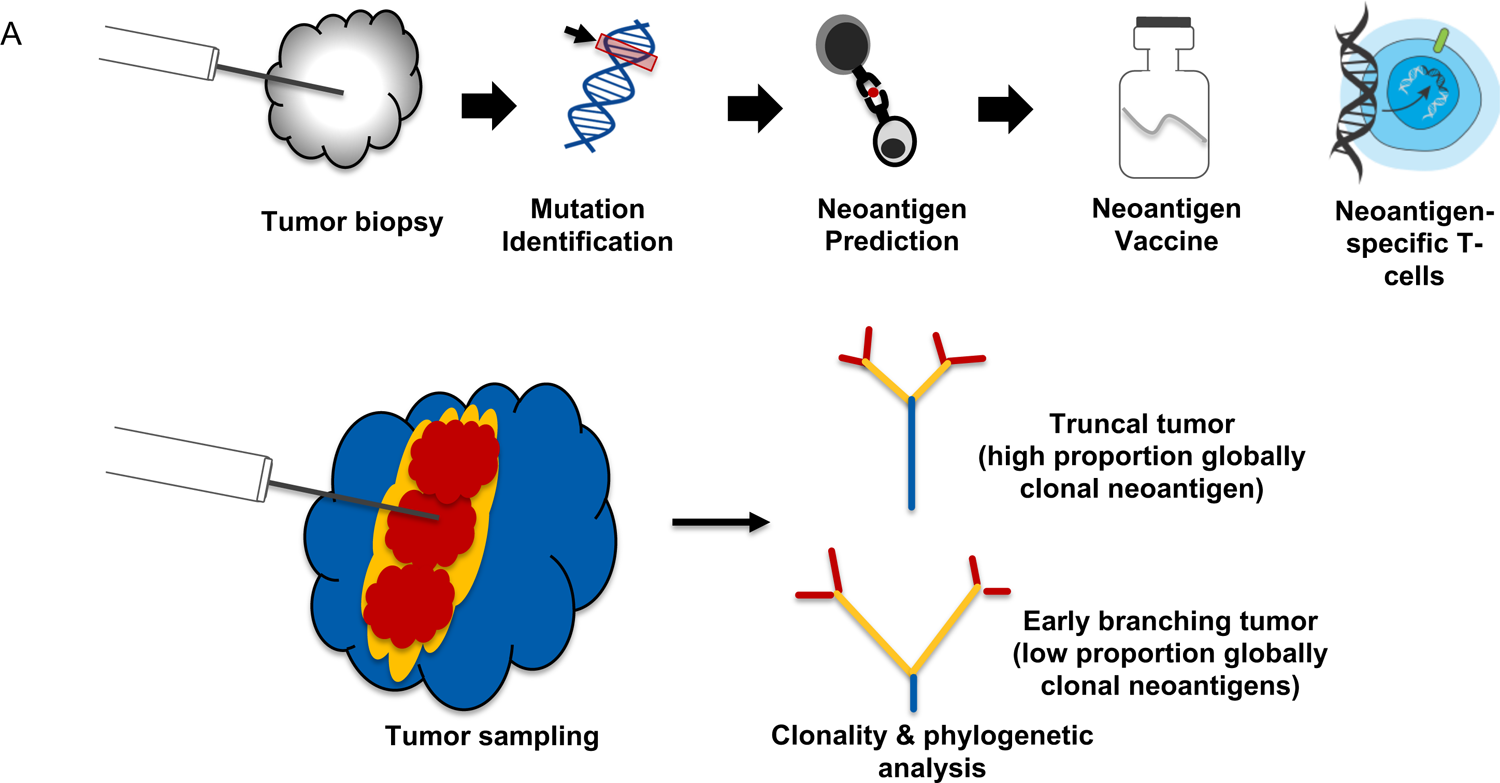
An example of an iNeST workflow, and neoantigen heterogeneity analysis by MR-Seq. (A) Individualized neoantigen-specific immunotherapy (iNeST) targets neoantigens, which are unique to an individual’s tumor. A neoantigen vacccine is an example of an iNeST. (B) Mutational heterogeneity in metastatic disease settings may pose a problem for iNeST targeting and efficacy. Mutation/neoantigen clonality varies across indications, with melanoma and NSCLC having low heterogeneity (highly clonal), and CRC, RCC, and breast cancer having high heterogeneity (low clonality). Predominantly primary tumors have been studied in non-metastatic disease setting, so the benefit of global mutation clonality prediction from standard clonality metrics determined using single tumor samples is unclear.

It has been suggested that clonal neoantigens should be targeted for neoantigen-specific immunotherapies to be most effective and to limit tumor escape, as clonal neoantigens are by definition present in every tumor cell ^6^. However, the clonal composition of metastatic solid tumors remains relatively under-explored, particularly in terms of neoantigen content across lesions. Various MR-seq studies have explored somatic mutation distributions across tumor lesions in patients with metastatic disease ^7–10^, but these studies are often complicated by samples being taken at different surgical timepoints as well as by intervening treatments. Various groups have attempted to create computational approaches to identifying clonal mutations from single biopsies. However, unambiguously identifying clonal mutations from single tumor biopsies is a challenge due to technical limitations on mutation detection (e.g., suboptimal tumor tissue quality, or stromal or immune infiltration). Identifying globally clonal mutations shared across tumor lesions presents yet another challenge in the clinic due to clonal heterogeneity within lesions and diverse processes underlying metastatic seeding (i.e., possible polyclonal tumor evolution) ^11^. Here we use the term “clonal” to refer to mutations inferred to be present in all tumor cells within a single sample or lesion, and the term “globally clonal” to refer to mutations empirically detected across tumor lesions regardless of their variant allele frequency (VAF). In contrast to targetable oncogene alterations such as those in *EGFR, ALK, PIK3CA, BRAF*, which are typically globally clonal ^12^, it is unclear how often private passenger mutations are shared across tumor lesions. Previous studies suggest that tumors with high heterogeneity due to branching evolution, where clones diverge from a common ancestor, should generally have a low proportion of globally clonal neoantigens ^13–15^, although tumors with high mutation loads may still have relatively high absolute numbers of clonal neoantigens. We therefore sought to characterize neoantigens in primary and metastatic tumors from four solid tumor indications taken at a single point in time to address the feasibility and implications of targeting globally clonal neoantigens.

Assuming that globally clonal neoantigens are preferable for individualized neoantigen specific immunotherapy (iNeST), several bioinformatics tools and approaches exist to estimate mutation clonality from tissue sequencing data. Empirical methods for calculating cancer cell fraction (CCF) rely on prior knowledge of genome-wide copy number alterations (CNA) from tumor/normal sequencing data, as well as the associated tumor purity estimates. These methods then normalize the VAF of each mutation to tumor purity and local copy number, with some using conditional probabilities to determine the exact adjustment needed on a mutation-by-mutation basis ^10, 11^. (We refer to the CCF estimates derived from empirical methods as “emp-CCF”). Separately, a number of computational tools have been developed that rely on Bayesian clustering of tumor VAFs, adjusted for local copy number and tumor purity, to derive CCF for each mutation (which we refer to as “clust-CCF”) ^16–18^. However, it remains unclear how well single-sample VAF, emp-CCF, or clust-CCF correlate with global mutation clonality across tumor lesions. We therefore investigated whether these mutation abundance metrics, as derived from single tumor samples, could predict global mutation clonality using an MR-seq-based “truth set” of mutations from each patient. We also explored whether single sample types (e.g., primary versus metastatic) had inherently better or worse predictive value for globally clonal mutations.

Levels of neoantigen presentation by MHC-I are correlated with neoantigen expression, and higher levels of presentation may trigger immune responses that subsequently lead to downregulation or removal by the tumor of the mutant allele underlying these immunogenic neoantigens. To generate efficacious iNeST, it is therefore important to understand the prevalence and characteristics of such “neoantigen depletion” by tumors, and how to consider this factor in neoantigen selection for iNeST. Neoantigen depletion can occur via genetic loss of mutations or loss of mutant allele expression, or by loss of MHC-I presentation, via loss of HLA alleles, as described in previous MR-seq studies ^14, 15^. Neoantigen loss through elimination in tumor subclones, chromosomal deletions and/or trucal alterations has been associated with changes in T-cell receptor clonality and could impact tumor response to immunotherapy ^19^. However, these various types of neoantigen depletion have not been systematically characterized in an unbiased fashion across lesions in metastatic tumors. We therefore extended the work of previous studies to explore whether the metastatic solid tumors in our study showed any evidence of neoantigen depletion via any of the above mechanisms. Taken together, our findings suggest that indication-specific tumor evolution, emp-CCF, and neoantigen depletion are important factors in neoantigen targeting, and that additional tumor sampling can help mitigate the limitations of single samples. Our findings have additional implications for the development of biomarker strategies for individualized cancer immunotherapies.

## Methods

### Clinical sample identification and pathology analysis

Institutional Review Board approval was obtained from Northwestern University. All surgical resection cases diagnosed as colorectal adenocarcinoma (CRC), non-small cell lung carcinoma (NSCLC), urothelial bladder carcinoma (UBC) or renal cell carcinoma (RCC) between 1992 and 2018. Three or more regions from primary tumor, three or more regions from lymph node metastasis close to the tumor and three or more regions from metastasis distant from the tumor as well as paired normal tissue from a single surgical time point were identified from the pathology database at Northwestern Memorial Hospital. All patient identifiers were reassigned to protect anonymity. A surgical pathologist reviewed all slides associated with each case, established that regions were located more than >1 cm from each other based on the gross description and confirmed diagnoses based on morphologic and immunohistochemical findings. Estimated viable tumor content (% viable tumor/total epithelial surface area), percent tumor nuclei (%viable tumor nuclei/total nuclei) and percent tumor area necrosis (% necrosis/total tumor area) for each case were then estimated by an independent pathologist. Retrospective chart review was performed to identify and capture relevant clinical and demographic information [Table 1. Cohort description and patient metadata].

**Table 1:**
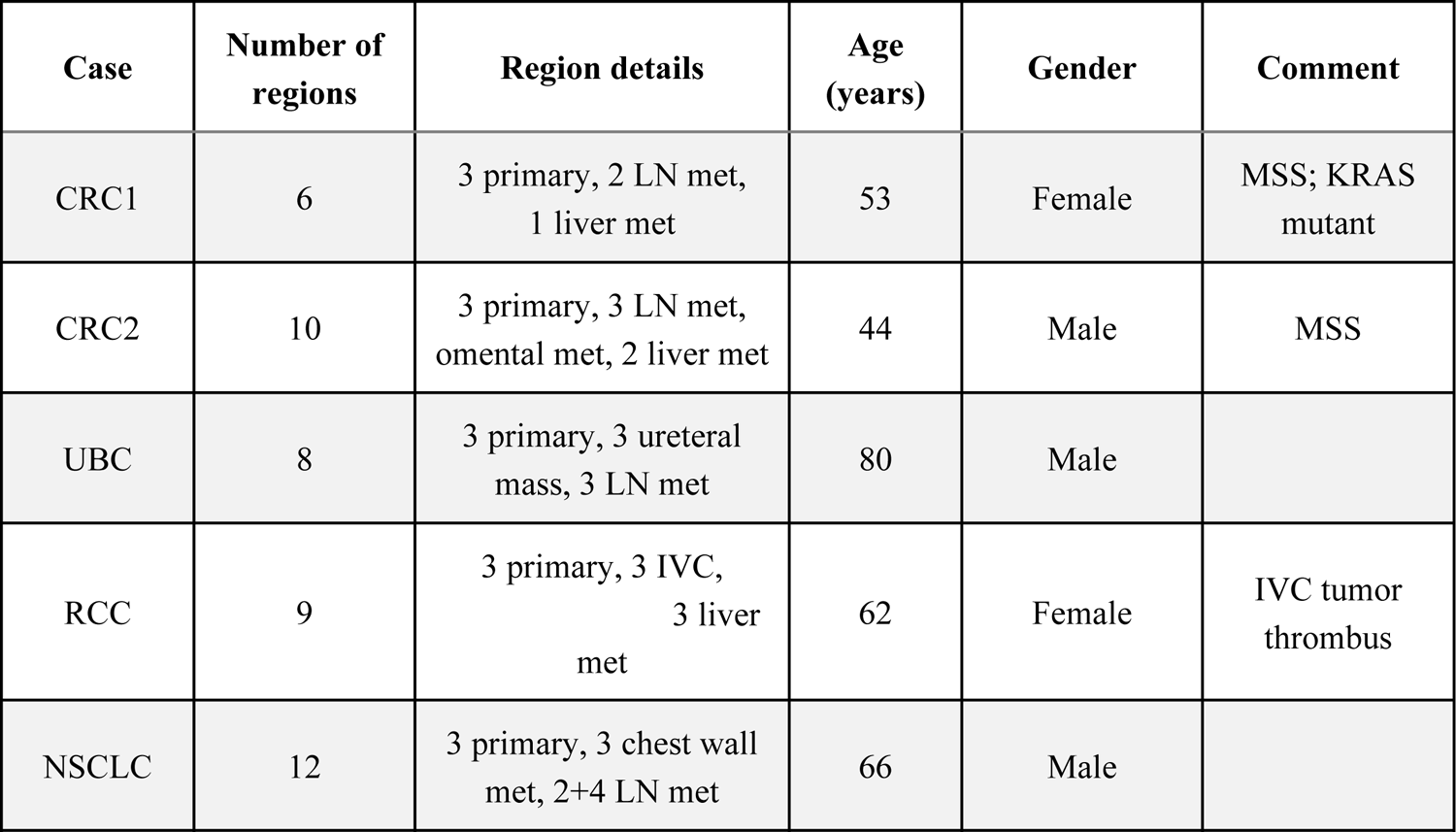
Clinical sample demographics for MR-Seq data generation. Case and indication breakdown with regions sequenced and patient demographics are shown. Multi-region exome and RNA sequencing, and IHC analysis, were performed on at least three regions from a primary tumor, three regions of a metastatic tumor, and matched normal tissue sampled at a single surgical time point.

| Case | Number of regions | Region details | Age (years) | Gender | Comment |
| --- | --- | --- | --- | --- | --- |
| CRC1 | 6 | 3 primary, 2 LN met, 1 liver met | 53 | Female | MSS; KRAS mutant |
| CRC2 | 10 | 3 primary, 3 LN met, omental met, 2 liver met | 44 | Male | MSS |
| UBC | 8 | 3 primary, 3 ureteral mass, 3 LN met | 80 | Male |  |
| RCC | 9 | 3 primary, 3 IVC, 3 liver met | 62 | Female | IVC tumor thrombus |
| NSCLC | 12 | 3 primary, 3 chest wall met, 2+4 LN met | 66 | Male |  |

### AVENIO Millisect tissue harvest

We applied rigorous quality control analysis to our tissue input by selecting cases with ischemic times less than 2 hours and maximizing viable tumor input using AVENIO Millisect automated dissection for tumor enrichment on all cases. Five or forty-nine cases demonstrated low tumor content (tumor areas below 30mm^2^) and were removed from downstream analyses. These were mostly lymph node metastases. A tissue processing, sequencing and data analysis workflow overview is presented in Supplementary fig. 1. Formalin Fixed Paraffin Embedded (FFPE) tissue blocks were serially sectioned with 5 sections at 10µm, followed by 5 sections at 4µm, collected onto Superfrost Plus positively charged slides (Thermo Scientific, Runcorn, UK) and allowed to dry at room temperature overnight. Serial section 6 (4µm) was baked at 60C for 30 minutes and stained with Hematoxylin and Eosin (H&E) on an automated Leica Autostainer XL using a routine protocol. H&E stained slides were scanned on a NanoZoomer 2.0 HT whole slide imager (Hamamatsu, Bridgewater NJ) at 20X magnification. Scanned slide images were annotated by a pathologist for tumor regions of interests and digital masks were created as a dissection reference.

Tissue sections were dissected using the reference mask image from serial section 6 to collect regions of interest using medium or large AVENIO Millisect milling tips (Roche Sequencing Solutions, Pleasanton, CA), collected with Molecular Grade Mineral Oil (Sigma-Aldrich, St. Louis, MO) as dissection fluid and dispensed into nuclease-free 1.5mL Eppendorf tubes. Dissections from slides 1 through 5 were centrifuged for 10 minutes at 20,000rpm to pellet tissue. Portions of mineral oil were removed from the tissue pellets and pellets were pooled in a single 1.5mL Eppendorf tube and held for DNA and RNA dual extraction. Post AVENIO Millisect dissected tissue slides were baked at 60C for 30 minutes and stained with Hematoxylin and Eosin (H&E) on an automated Leica Autostainer XL using routine protocols and scanned on a NanoZoomer 2.0 HT whole slide imager (Hamamatsu, Bridgewater NJ) at 20X magnification in order to confirm that selected tumor regions were successfully removed from the slides. DNA and RNA extraction was performed using the Qiagen AllPrep DNA/RNA tissue kit (Qiagen, Germantown, MD) at Q^2^ Solutions (Valencia, CA).

Tumor content ranged from 1 to 90% in tissue regions analyzed and tumor enrichment was performed on all cases using AVENIO Millisect for semi-automated dissection resulting in tumor input of 2.5-1950mm^2^ (Supplementary Table 1) that excluded any surrounding normal tissue and necrotic regions from capture and analysis (Supplementary fig. 2). Cases with input less than 30mm^2^ were removed from the analysis. Matched normal tissue was dissected from separate tissue blocks in regions >5cm away from any tumor mass for DNA extraction.

**Fig. 2:**
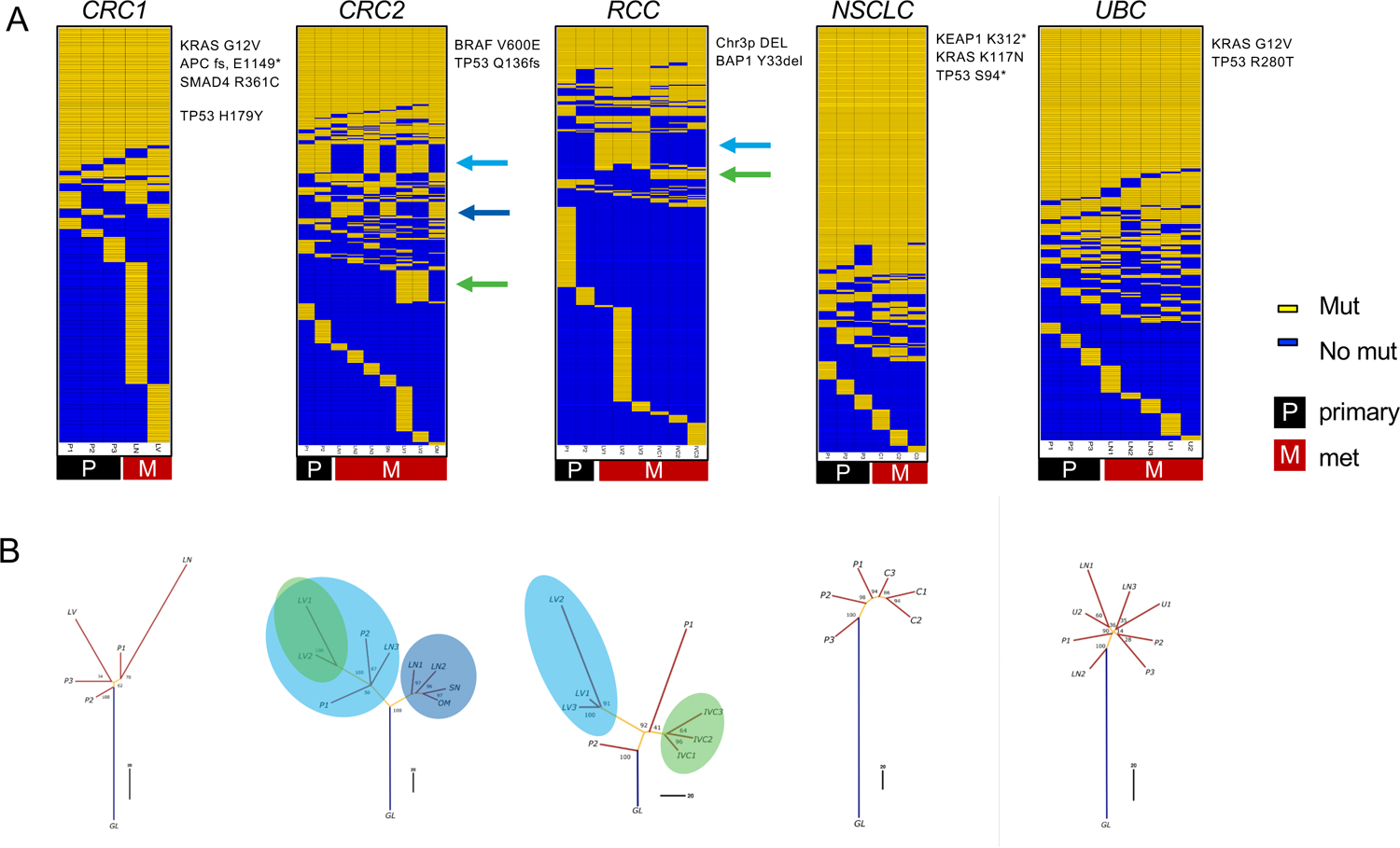
Evolutionary analysis reveals clade-specific neoantigens in CRC2 and RCC cases. (A) Somatic mutation calls were used to create sorted binarized mutation heatmaps for each case. (B) The mutation data were also used to reconstruct mutational phylogenies of each tumor. The early branching evolutionary pattern in RCC and CRC2 cases contrasts with a more truncal pattern in the NSCLC and UBC cases. For the RCC and CRC2 cases, the green and blue arrows in (A) and circles in (B) indicate clade-specific mutations shared across multiple tumor regions. P = primary, LN = lymph node, LV = liver met, OM = omental met, SN = satellite nodule, C = chest wall met, IV = tumor thrombus, U = uretal mass, GL = germline.

### Multi-region sequencing and exome data analysis

The quantity of isolated DNA and RNA was determined using a Qubit, and the quality and fragment lengths were confirmed using a BioAnalyzer. DNA sequencing libraries were created with Agilent SureSelectXT and libraries were used for hybridization and capture with SureSelect All Exon V6 bait probes at Q^2^ Solutions (Valencia, CA). Whole-exome sequencing (WES) coverage was approximately 75 million 100bp paired-end reads, yielding an average depth (before removing duplicate reads) of 150X per sample. RNA sequencing (RNA-seq) libraries were generated using the RNA Access platform. Sequencing coverage was approximately 50 million 50bp paired-end reads per sample. DNA and RNA sequence read alignments were performed using *GSNAP* ^20^, which was run within the *HTSeqGenie* pipeline for sequence read alignment and QC (see https://bioconductor.org/packages/release/bioc/html/HTSeqGenie.html). Full alignment statistics for RNA-seq can be found in Supplementary Table 5.

Somatic mutation calling was performed using *Strelka* v1.0.14 ^21^ and *LoFreq* ^22^, and a combined VCF file containing the union of calls from the two callers was generated for each sample. Somatic mutation calls were carefully filtered using read coverage, VAF, allele frequency in normal sample (NAF), and ExAC (Exome Aggregation Consortium) GMAF (global minor allele frequency) criteria such that only high confidence mutations were included in downstream analyses. To be included in downstream analysis, a mutation had to meet the following criteria: VAF >= 0.05 in at least one sample, a coverage minimum of 20 reads in at least one sample, a maximum ExAC GMAF of 0.01, and a maximum NAF of 0.01.

Mutational signatures were generated using the *MutationalPatterns* R package ^23^. Clonal copy number analysis was performed using *TitanCNA* ^24^. Identification of putative neoantigens was performed using custom code to annotate and translate in silico transcripts containing the mutations to mutant peptide sequences. HLA class I alleles were called from the matched normal exome sequencing data for each patient using *HLA-HD* ^25^. Neoepitope presentation was then predicted for tumor-specific peptides of length 8-11 using the eluted-ligand mode of *NetMHCpan-4.0* ^26^.

Sample and sequencing data quality control was performed using several metrics from the sequencing alignments, somatic mutation calls, and copy number calls. For each sample, the number of uniquely mapping reads and cumulative coverage distribution were first examined, followed by VAF distributions, and finally genome-wide logR signal for somatic copy number. Tumor samples that had compressed VAF distribution (median VAF < 0.1 and IQR < 0.5) or low median logR values when visualized in IGV were removed from the analysis. These QC procedures led to the removal of four lymph node samples from the NSCLC case, as well as one primary tumor region from the CRC2 and RCC cases. All downstream analyses were performed using custom scripts in R.

#### Phylogenetic analysis

Mutation matrices were constructed for each patient by joining together the VAF values for the mutations called in each tumor region. The VAF values were binarized to a discrete character set (0=absence and 1=presence of the mutation in the sample), and the binarized mutation matrices were used to plot heatmaps and to create mutational trees (fig. 2). The full set of input mutations passing the previously described filters was used. An additional “GL” (germline) sample was introduced as an out-group containing all zeroes, setting the ancestral state of all the tumor mutations. Tumor mutational phylogenies were constructed with the R package *phangorn* ^27^. For each patient, a maximum parsimony tree was generated using the parsimony ratchet method ^28^ implemented in the function phangorn::pratchet(). Branch lengths were determined by the Hamming distance between all the samples involved in a tree as an input to the non-negative least squares method implemented in the function phangorn::nnls.phylo(). Finally, bootstrapping to estimate the confidence of the tree topology values was performed by re-sampling 100 trees from the data using the function ape::boot.phylo() from the *ape* R package ^29^. Tree plotting was then performed using standard R functions.

#### Global clonality analysis and VAF/CCF comparisons

A mutation was called “global” if it was found in all regions of a given tumor. The *percent global* for mutations in each tumor sample was calculated as:

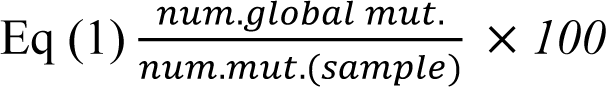

In contrast, the *total global fraction* (equivalent to the percentage of all unique mutations in a tumor that are shared across all tumor regions) was calculated as:

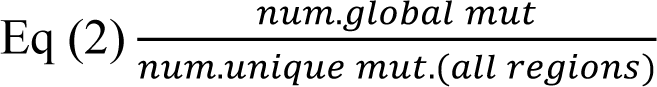

*VAF* was calculated using the standard approach per mutant site:

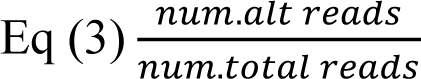

*emp-CCF* was calculated following the method of Turajlic et al. ^10^:

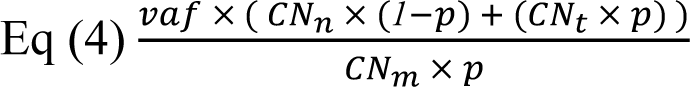

where *p* is estimated tumor purity from TitanCNA, *CN_n_* is the total copy number for the overlapping segment in the matched normal sample (assumed to be 2 in all cases), *CN_t_* is the total copy number for the overlapping segment in the tumor sample, and *CN_m_* is the copy number of the mutant allele. An important exception was that *CN_m_* was assigned a fixed integer value for each mutation, using the MajorCN value for the overlapping segment from TitanCNA. In calculating emp-CCF, there were approximately 100 mutations across the five cases where the tumor purity integer CNA values appeared incorrect, and there was no clear way to determine the correct integer copy number (these mutations were removed from the analysis). Additionally, there were approximately 50 mutations where VAF, local copy number, and/or tumor purity were incorrect and the CCF values were slightly greater than 1 (we adjusted these to emp-CCF=1).

*Clust-CCF* was determined by providing the combined somatic mutation calls (union) from Strelka and LoFreq and the CNA calls from TitanCNA to *phyloWGS v1.0-rc2* ^18^. PhyloWGS was run with standard parameters, including 1,000 burn-in samples and 2,500 MCMC samples. The highest likelihood tree was then taken, and the resulting *phi* values for each cluster were converted to *clust-CCF* values using *phi/max(phi)*.

### Neoantigen depletion analysis and RNA expression signatures

Quantification of neoantigenic allele expression was performed using custom R and Python code to count variant and reference allele-containing read pairs. The Python code made use of pysam (https://github.com/pysam-developers/pysam), which wraps samtools ^30^. The ratio of variant-containing read pairs to total read pairs was multiplied by the gene-level RPKM to estimate a variant-containing RPKM. To assess neoantigen expression as a function of neoantigen presentation, the variant allele counts, as well as the variant RPKM values, were compared for neoantigens with EL_mut_ <= 2 (presented) and those with EL_mut_ > 2, using Mann-Whitney U tests. Where appropriate, correction for multiple testing was performed using the method of Benjamini and Hochberg ^31^.

RNA-seq data was used to estimate the relative infiltration of B cells, dendritic cells, macrophages, neutrophils, NK cells, CD4 T cells, CD8 T cells, CD8 T-effector cells, Th1 cells, and Th2 cells in tumor samples. Gene expression signatures were derived for these cell types using the Danaher et al. method ^32^. The mean cross-sample-normalized expression values of cell type signature genes was then used as a proxy for the relative infiltration of each cell type. The correlation of the CD8 T cell signature and the IHC CD8 density estimates was then assessed.

### IHC analysis

Immunohistochemistry (IHC) was performed on 4um thick formalin-fixed, paraffin-embedded tissue sections mounted on glass slides. IHC for PD-L1 clone SP263 (Roche Tissue Diagnostics, Tuscan, AZ, cat 790-4905) was performed on the Ventana Benchmark XT platform. The slides were pretreated with CC1 for 64 min followed by primary antibody incubated for 16 minutes at 37C. The antibody was detected with the OptiView DAB IHC Detection Kit.

PanCK (vendor) and CD8 (vendor) duplex chromogenic IHC was performed using established methods on the Ventana Discovery Ultra. The fraction of viable tumor cells (%) that express membrane PD-L1 were quantified. The overall immune phenotype was classified as desert, inflamed, excluded based on the predominant (>10%) location of CD8 positive cells in relation to the tumor. Automated slide assessment was performed quantitatively using Visiopharm analysis software. Tissue area that was positive for the panCK was used to generate an epithelial tumor mask and the relative surface area of CD8 positive cells within stromal and epithelial tumor compartments was determined using Visiopharm software applications. CD8 density as absent (0), low (1), moderate (2) or high (3) in intratumoral panCK positive areas and intratumoral panCK negative (stromal) areas was captured and calculated as an H-score.

## Results

### MR-seq analysis reveals distinct evolutionary modes across indications and clade-specific neoantigens

We reasoned that characterizing mutation presence/absence and mutation expression across tumor regions could yield insights into how neoantigens and their related properties are distributed across lesions in metastatic cancer patients. We therefore used MR-seq to identify somatic mutations (WES), to determine their expression levels (RNA-seq), and to predict neoantigens from the *in silico* translated mutant peptide sequences across tumor regions. We sequenced 42 tumor regions across 5 patients, including 15 primary regions and 27 regions from metastases, as well as a matched normal sample distant from the tumor from each patient (Table 1). Following sample quality control procedures, 36 tumor regions with WES and 35 tumor regions with RNA-seq were used for downstream analyses. Somatic mutation analysis revealed the expected patterns of base substitution and mutational signatures previously established for each indication (Supplementary fig. 3) ^33, 34, 35–37^. Several known driver mutations and CNAs commonly associated with each indication were identified in each tumor (Fig. 2A, Supplementary fig. 4), and these alterations were generally globally present across tumor samples.

Phylogenetic analysis of binarized somatic mutations from each tumor identified striking cross-indication differences in how mutations were or were not shared across tumor lesions. While the term “truncal” can be used to describe individual mutations or overall phylogenies, we use it to describe a mode of tumor evolution in which a single clone grows out and persists through metastasis. In contrast, “branching” refers to a mode of evolution in which multiple mutationally distinct tumor clones grow out early in tumor development. (See Fig. 1B.) The CRC2 and RCC cases followed a branching evolutionary mode, with low proportions of globally clonal mutations (Fig. 2A and B). The early branching in these cases led to a relative dearth of globally clonal neoantigens, with 15 in CRC2 and nine in RCC. In contrast, the NSCLC and UBC cases followed a predominantly truncal evolutionary mode, with higher proportions and absolute numbers of globally clonal mutations (Supplementary Table 2). The CRC1 case appeared intermediate between these two groups of cases. These results are generally consistent with previous findings in these indications ^9,38, 39^.

Notably, the CRC2 and RCC cases each harbored sets of clade-specific shared mutations (or in evolutionary terms, a “synapomorphies”) that were shared by distinct tumor lesions and regions. In the RCC case, there were 36 clade-specific mutations (10 clade-specific neoantigens): 28 mutations (7 neoantigens) were exclusive to the liver met, and 8 mutations (3 neoantigens) were exclusive to the IVC met. In the CRC2 case, there were 74 clade-specific mutations (15 clade-specific neoantigens): 14 mutations (3 neoantigens) were exclusive to the LN1, LN2, OM, and SN mets, 28 mutations (5 neoantigens) were exclusive to the primary regions, LN3 and liver met, and an additional 32 mutations (7 neoantigens) were exclusive to the liver met. Thus, considering clade-specific neoantigens in addition to globally clonal neoantigens would effectively double the total set of neoantigens shared across tumor regions in both of these cases.

### Globally clonal neoantigen numbers and proportions vary consistently across indications

MR-seq provides an ability to measure both the total numbers and proportions of globally clonal neoantigens across indications. We reanalyzed 103 published MR-seq cases from several studies to determine how the abundance of global mutations varied across the same four indications ^9, 38, 40–43^. We found that the total numbers of global mutations varied nearly seven-fold across indications, with RCC on the low end (median, 26) and NSCLC on the high end (median, 173) (Fig. 3). We then inferred which of these mutations would represent likely neoantigens, and found that the median number of global neoantigens in these indications ranged from 12.8 in RCC to 86.5 in NSCLC. CRC and UBC fell in between, with UBC tumors having more global mutations and neoantigens and CRC tumors having fewer. A caveat of this analysis is that we were not able to match disease stage across indications, with NSCLC in particular having almost no published metastatic MR-seq cases.

**Fig. 3:**
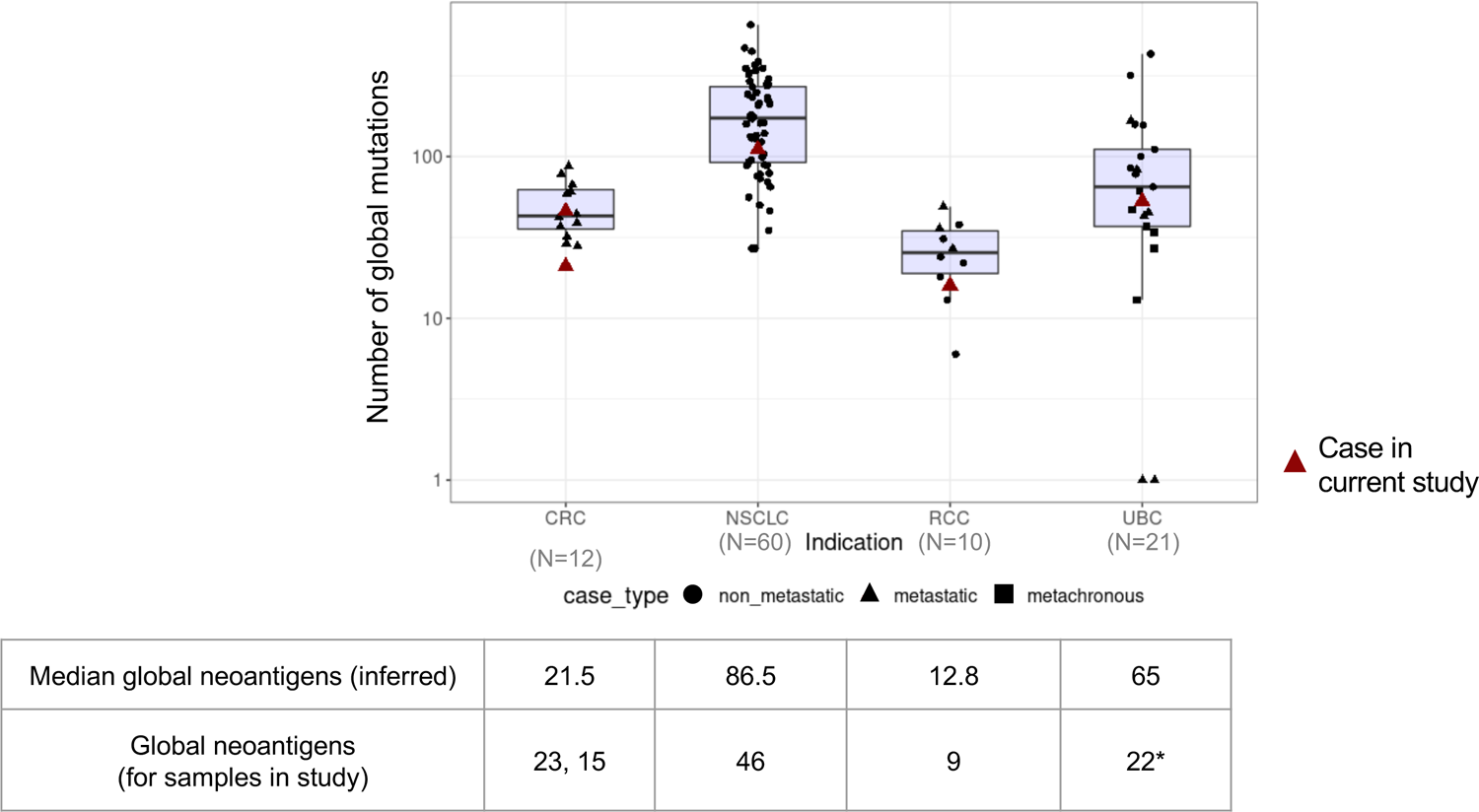
Absolute numbers of global mutations across indications. The absolute numbers of global mutations per tumor were determined from published MR-seq studies. The analysis was limited to cases with at least three tumor regions. The five cases from the present study are indicated by red triangles. Note that all published NSCLC were non-metastatic and there was some intervening treatment in the UBC studies. Mutation data were collected from Hu Nat Gen 2019, Jamal-Hanjani NEJM 2017, Gerlinger Nat Gen 2014, Lamy Cancer Res 2016, Faltas Nat Gen 2016, Heide J Pathol 2019. Median global neoantigens were inferred by multiplying the observed ratio of neoantigens to mutations in the current study by the median number of mutations in the published studies, in an indication-specific fashion. *The number of global neoantigens in the UBC case in the present study is lower than expected due to many neoantigenic mutations not being expressed (see Supplementary Table 2 and Fig. 6).

### Proportions of globally clonal mutations differ when sampling single primary versus single metastatic tumor sites

Because most neoantigen-specific immunotherapies rely on up-front identification of somatic mutations from tumor sequencing data, we sought to understand how well a given tumor sample could capture globally clonal mutations. To address this, we considered the set of mutations found in all samples from a given patient to represent the globally clonal set and then asked what percentage of the mutations found in a given tumor sample were in that set (Fig. 4A, see Eq. 1 in Methods). We note that the percentage of mutations that are global per sample (“percent global”) is equivalent to the predictive value of that tumor sample for the identification of globally clonal mutations. Thus, a tumor sample with 60% global mutations would yield a ∼60% probability of identifying a global mutation when choosing randomly. Samples with a higher percentage of global mutations should therefore have an inherently higher likelihood of yielding global neoantigen targets for immunotherapy.

**Fig. 4:**
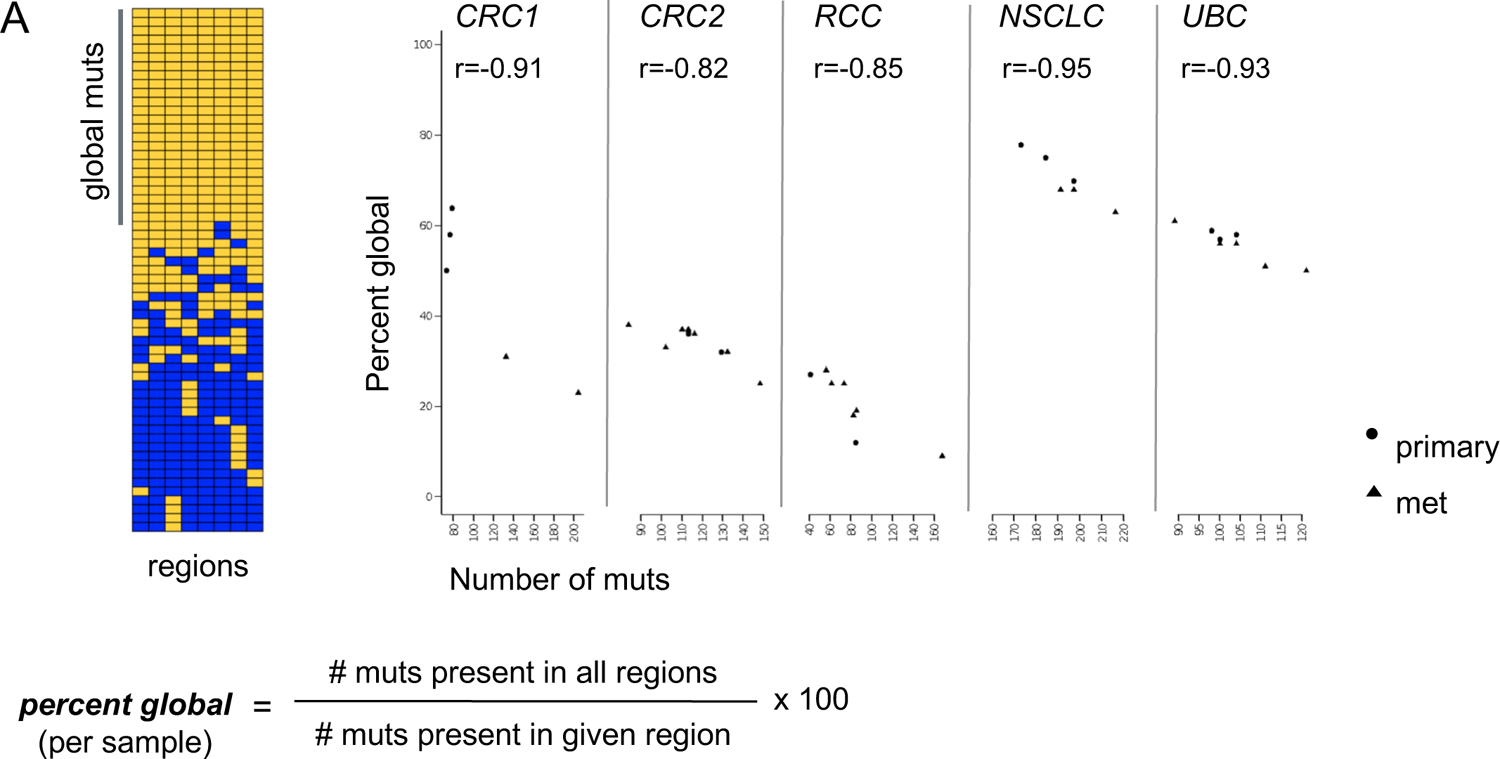
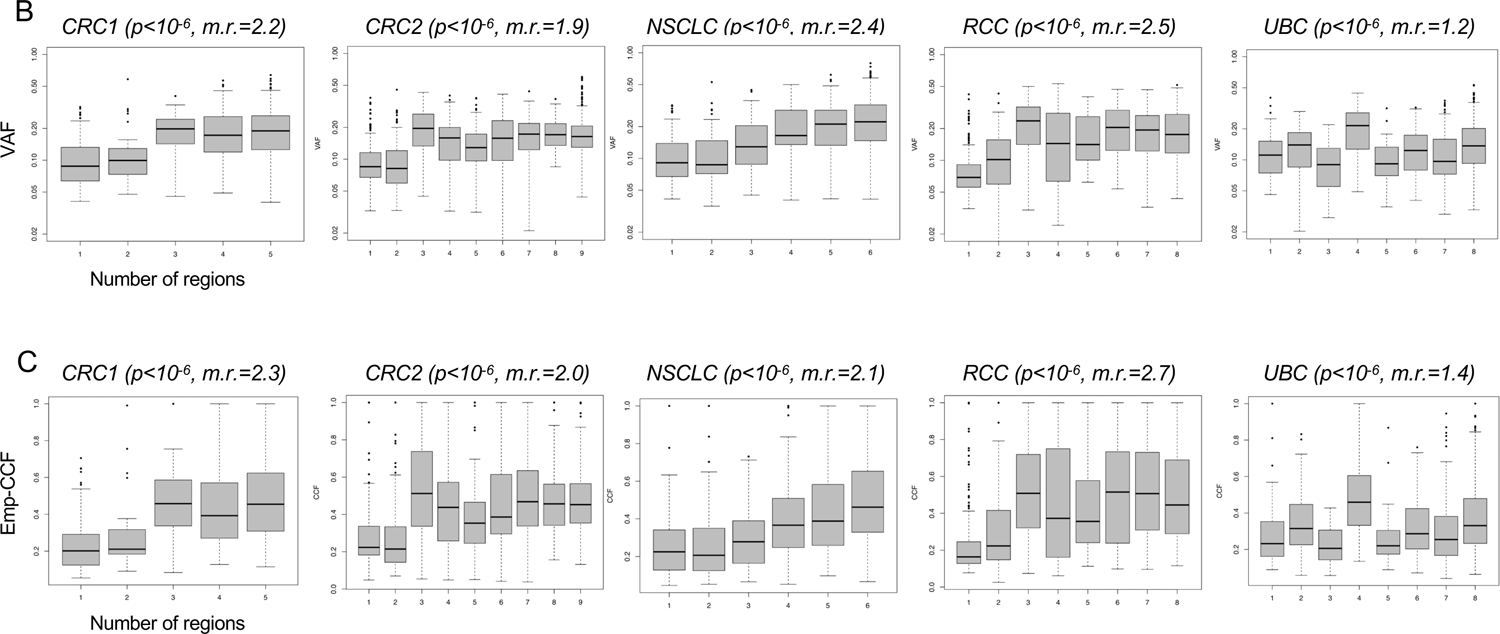
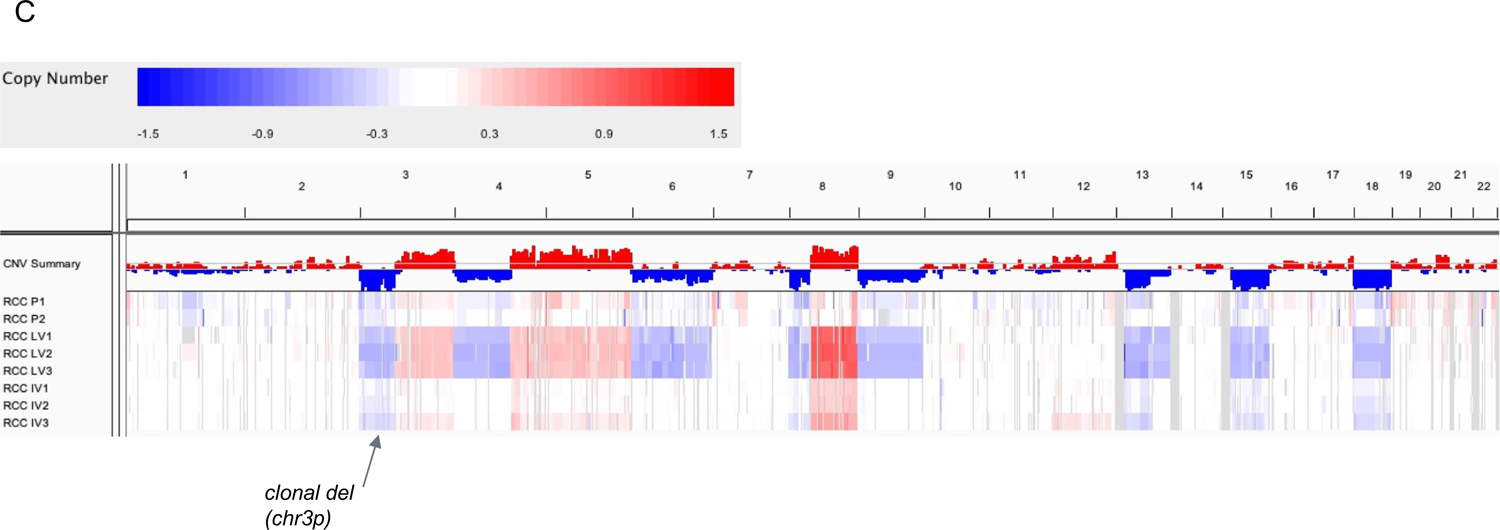
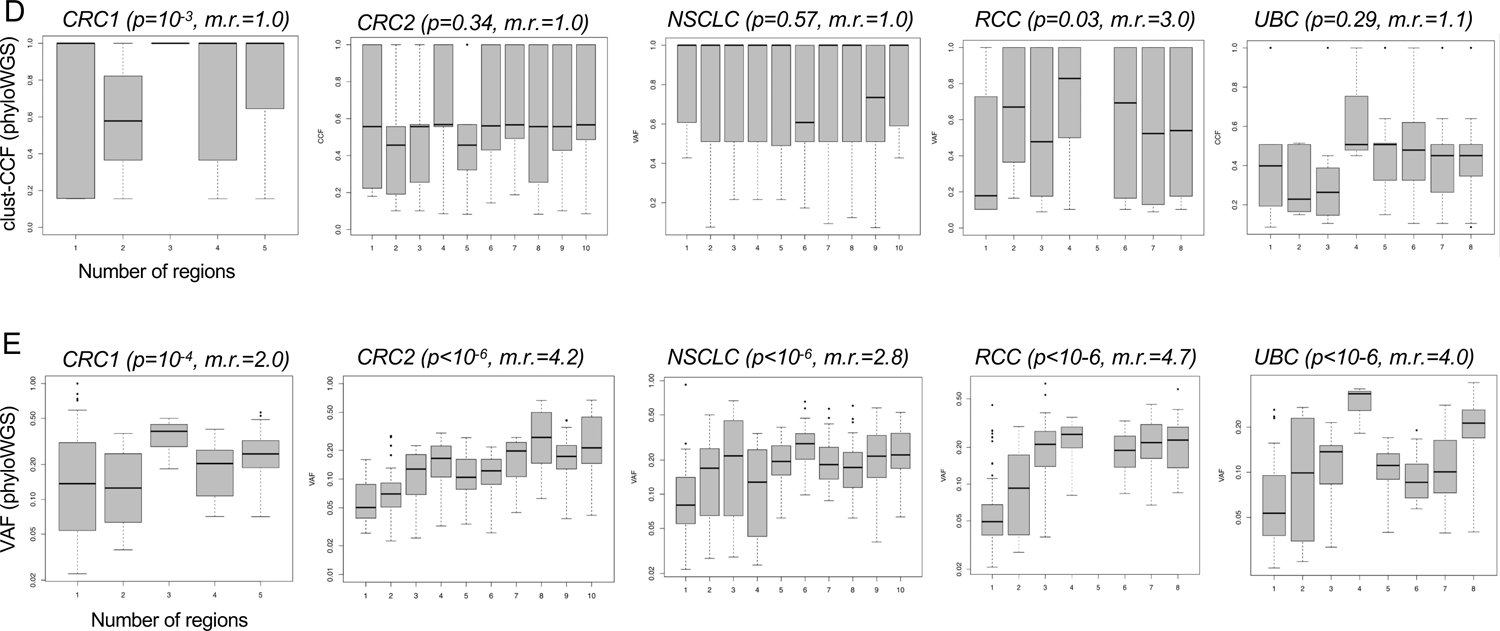
Predictive value of individual tumor samples for globally clonal mutations, and comparison of VAF, emp-CCF, and clust-CCF with mutation presence across tumor regions. (A) MR-Seq can inform tissue sampling for iNeST. The “percent global” statistic can also be used as a measure of predictive value for each tumor region for globally clonal mutations (those found in all regions). Primary samples generally had higher predictive value for clonal mutations than metastasis samples did. Mets tended to have more mutations (p=0.03), consistent with later clone emergence. Some lymph node samples suffered from severe under-detection of mutations due to inadequate tumor tissue area, and were removed from all analyses (see Methods). (B) Variant allele frequencies (VAF) are plotted for mutations found in different numbers of regions for each case. The VAF distribution for singleton mutations (those found in just one region) are at the left-most x-axis position, and the VAF distribution for globally clonal mutations (those found in all regions) are at the right-most x-axis position of each box plot as the number of regions increases from left to right on the x-axis. The p-value from a Kruskal-Wallis test (VAF ∼ num_regions) is indicated in parentheses next to each case name, as well as the ratio of median VAF for clonal mutations divided by the median VAF for singleton mutations (“m.r.”). (C) Empirical cancer cell fractions (emp-CCF), as calculated using Equation 4 (see Methods) are plotted for mutations found in different numbers of regions for each case. Results from similar analyses as performed in (B) are shown, with p-values from Kruskal-Wallis tests (emp-CCF ∼ num_regions). (D) Cancer cell fractions (CCF) determined by phyloWGS are plotted for mutations found in different numbers of regions for each case. Results from similar analyses as performed in (B) and (C) are shown, with p-values from Kruskal-Wallis tests (clust-CCF ∼ num_regions) and the ratio of median clust-CCF for global mutations divided by the median VAF for singleton mutations (“m.r.”). (E) Variant allele frequencies (VAF) from the set of mutations passed to phyloWGS are plotted for mutations found in different numbers of regions for each case.

The percent global for each sample tended to vary across samples and patients and was also associated with indication (Fig. 4A). Across the five cases, the median percent global varied from 20% in the RCC case to 70% in the NSCLC case. Percent global was also negatively correlated with the number of mutations per sample, suggesting that samples with fewer mutations may yield a higher proportion of global neoantigens on average. In the CRC1 and NSCLC cases, primary samples tended to have fewer mutations and higher percent global, whereas metastasis samples tended to have more mutations and lower percent global. Although this trend did not clearly hold across the other cases, it was notable that all but one primary sample had at or above the median percent global mutations across all five cases. Finally, the lymph node metastases in the NSCLC case likely suffered from severe mutation under-detection due to inadequate tumor input secondary to low tumor area in the sample (Supplementary fig. 2 and Supplementary Table 1).

It has been suggested that sequencing a second tumor sample can help enrich global mutations ^44^. We tested how consistently second samples would enrich for global mutations using mutation set analysis (Supplementary fig. 5). We found that although a second sample would help at least modestly in all five cases, it led to the best enrichment of global mutations in the CRC1 case, which had numerous singleton mutations, or mutations found in only one tumor region, in some samples. In the CRC2 and RCC cases that demonstrated a moderate number of singleton mutations, sequencing a second sample would enrich both global mutations and clade-specific mutations. The NSCLC case harbored relatively few singleton mutations across samples, and a second sample would have only marginal benefit. Sequencing an additional tumor sample in some indications could therefore help, but does not readily allow one to distinguish clade-specific mutations from global mutations.

### VAF and emp-CCF, but not clust-CCF, can enrich for globally clonal neoantigens in single tumor samples

Having established the percent global mutations found in each sample, we next asked whether standard mutation abundance metrics, as determined from single samples, could be used to enrich for global mutations. We compared single-sample VAF to the number of regions in which a mutation was present. As expected, higher VAF was significantly associated with mutation presence in multiple regions across all five cases (Fig. 4B). Global mutations had median VAF ranging 1.2-2.5-fold higher than that of singleton mutations, suggesting that VAF alone could enrich global mutations in single tumor samples. We next compared emp-CCF to the number of regions in which a mutation was present. Higher emp-CCF was significantly associated with mutation presence in multiple regions across all five cases, and global mutations had median emp-CCF ranging 1.4-2.7-fold higher than that of singleton mutations (Fig. 4C). This suggested that emp-CCF may be marginally better at enriching for singleton mutations than VAF, but with important technical caveats related to tumor purity and local copy number estimation (see Methods).

We next compared clust-CCF to the number of regions in which a mutation was present. Surprisingly, clust-CCF values were poorly associated with mutation presence in multiple regions, and median clust-CCF was not substantially higher among global mutations compared with singletons except in the RCC case (Fig. 4D). Although many somatic mutations had to be dropped from this analysis due to the requirement for overlapping copy number segments in all tumor regions (see Methods), we note that this did not explain the poor association of clust-CCF with global mutation clonality, as VAF from the same subset of mutations was still robustly associated with global mutation presence across all five cases (Fig. 4E).

Additionally, we noticed that the presence of even minimal mutant allele expression significantly enriched for globally clonal mutations (Supplementary Table 2). Across all five cases, comparisons of the total global fraction (see Eq. 2 in Methods) for expressed mutations versus for all mutations suggested that the presence of mutant allele expression in a tumor sample provided a robust enrichment of global mutations (CRC1: OR=3.7, p=10^-5^; CRC2: OR=2.3, p=5×10^-4^; NSCLC: OR=1.8, p=2×10^-3^; RCC: OR=3.0, p=9×10^-4^; UBC: OR=2.5, p=10^-4^ by Fisher’s exact test).

### Evidence for PD-L1 and CD8 heterogeneity

It has long been recognized that levels of CD8+ T cell infiltration and PD-L1 expression within the tumor microenvironment provide important information regarding patient prognosis and likelihood of response to treatment. While thorough investigations into each of these biomarkers across several indications has been done^45, 46^, to our knowledge, analysis of differences across primary and metastatic regions at a single time point has not been performed. We therefore investigated tumor CD8 levels and CD8+ T cell localization as well as PD-L1 tumor and immune cell expression by immunohistochemistry. We detected significant PD-L1 tumor and immune cell expression heterogeneity across both primary and metastatic regions and in some tumors in multiple indications (Fig. 5C, D). In the UBC case in particular, regional PD-L1 variability was present and would have impacted histopathologic classfication based on standard scoring algorithms for UBC, an indication where scoring could impact treatment decisions; however, only one region for this case (a lymph node metastasis with less than 25% tumor cell PD-L1 expression) would have produced scores leading to a different treatment decision for the patient. Intraepithelial and intrastromal CD8 density was also heterogeneous across regions, but primary or metastatic regions tended to demonstrate similar levels of intraepithelial or intrastromal CD8 cell infiltration in some cases (Fig. 5A, B). Interestingly in the RCC case, CD8 intraepithelial and intrastromal content was higher in regions of tumor thrombus and metastatic regions compared to primary regions, which could support branching evolution in the development of these distinct regions or represent less well-established immunosuppression (Fig 5A, B). Estimated neoantigen load was not significantly correlated to tumor intraepithelial CD8 levels assessed by H-score (Fig. 5E), with the RCC case being a possible exception.

**Fig. 5:**
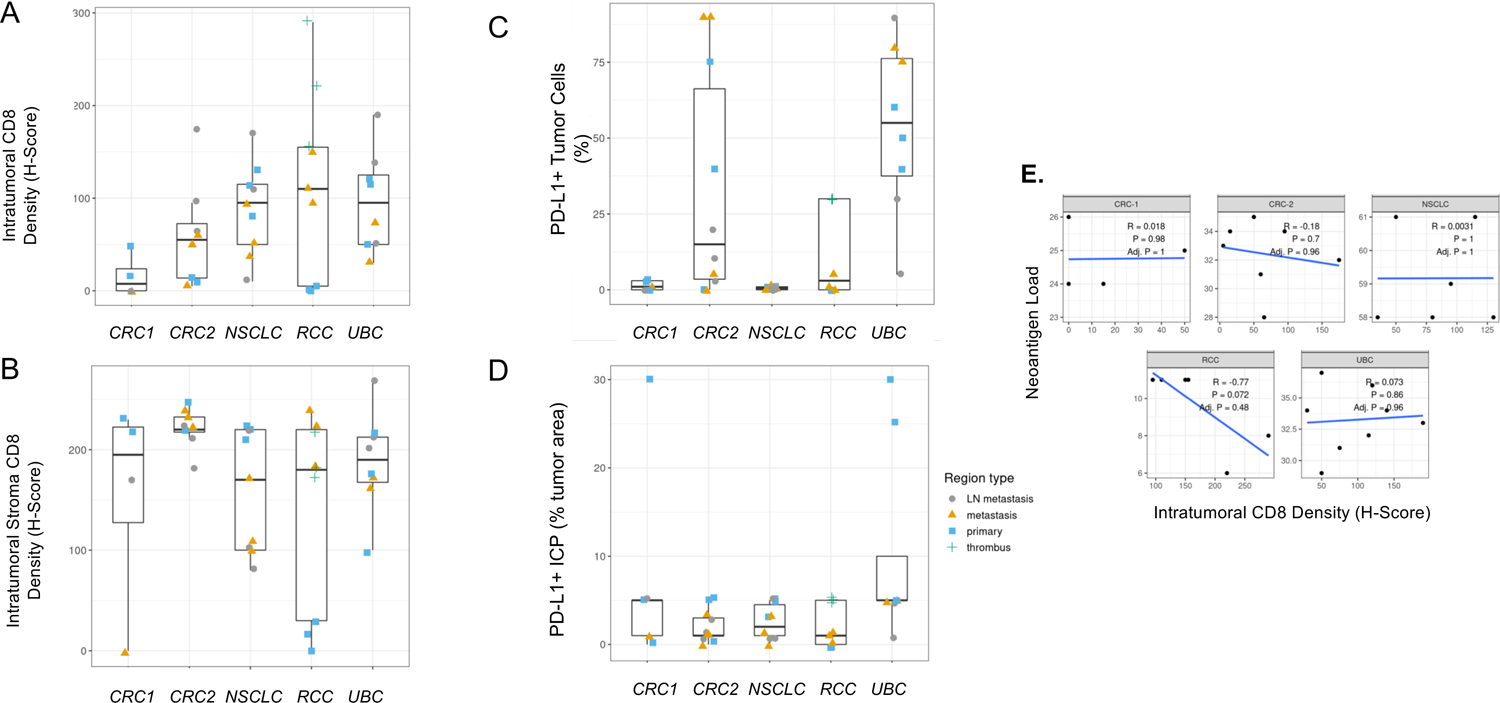
PD-L1 and CD8 IHC heterogeneity across indications. (A) Intratumoral CD8 T-cell density and (B) intratumoral stroma CD8 T-cell density across regions of each tumor by IHC H-score demonstrate that heterogeneity across regions may influence interpretations from single region analysis. (C) PD-L1 positive tumor cells across cases by IHC (SP263 clone) show that region-to-region variation in PD-L1 IHC signal can impact interpretation of immune phenotype for a tumor, but this would have impacted a treatment decision in only one UBC region that shows less than 25% of tumor cells positive. (D) The percent of tumor area occupied by PD-L1 positive tumor associated immune cells (immune cells present, ICP) across cases were generally less heterogeneous but demonstrated large differences in a few regions. (E) Neoantigen load correlation to tumor intraepithelial CD8 levels assessed by H-scores across regions for each case lacked statistical significance.

### Neoantigen depletion in NSCLC, RCC, and UBC cases occurs via distinct mechanisms

Previous studies have suggested that neoantigens can become depleted in untreated tumors via genomic deletion or expression down-regulation to enable tumor escape following immune recognition ^15^. We systematically examined the five metastatic cases for evidence of neoantigen depletion by mutation loss via copy number alterations (genomic deletion or LOH spanning the mutation), loss of mutant allele expression, and genetic loss of class I HLA alleles. We first looked for neoantigenic mutations (those giving rise to peptides with minimum *EL_mut_* scores <2) with overlapping copy number loss alterations in multiple tumor regions. We required that either the mutation VAF be reduced in regions harboring the CNA loss, or that the CNA be in mutually exclusive tumor regions relative to the neoantigenic mutation. Overall there were relatively few instances of genetic neoantigen loss, although at least one neoantigenic mutation found in one tumor region of the NSCLC primary was lost from a metastasis region as well as other primary regions due to copy number loss (Supplemental Table 3). Similarly there were a few neoantigens that were present in the RCC primary tumor but lost in all regions of the liver metastasis. These apparent neoantigen losses represented a small proportion of the total neoantigens in these cases, and overall neoantigenic mutations were not enriched in regions of CNA loss relative to all nonsynonymous mutations. This observation held true whether RNA-seq support for the alternative allele was disregarded (Supplementary Table 3A), or required (Supplementary Table 7A).

We next looked for expression depletion of neoantigens using two approaches. We first looked for variable mutant allele expression across regions of each tumor (lower mutant allele expression for mutations encoding neoantigens in a subset of tumor regions), and then asked if there was a clear trend where non-primary regions had significantly lower expression than primary regions. We did not find substantial evidence for neoantigen expression depletion using this approach. We next looked for association of mutant allele expression with neoantigenic status, and we found a significant trend in the UBC case where neoantigenic mutations had consistently lower expression across tumor regions than non-neoantigenic mutations (Fig. 6A and B). This trend could be observed either in aggregate or when looking across individual UBC tumor regions, and was statistically significant in three of eight regions (Supplementary fig. 6A). One of eight regions retained statistical significance after correction for multiple testing (Supplementary Table 6A). With use of raw alternative allele-supporting read counts as an alternative metric, 7/8 of the comparisons were significant after multiple testing correction (Supplementary fig. 6B, Supplementary Table 6B). These results suggested that the UBC tumors may have employed neoantigen expression depletion as a mechanism to evade immune surveillance. We identified 14 candidate neoantigens that were shared across tumor regions and appeared to mediate the depletion effect, as when these neoantigens were removed the trend largely disappeared in all tumor regions (Supplementary Table 3, Supplementary fig. 7).

**Fig. 6:**
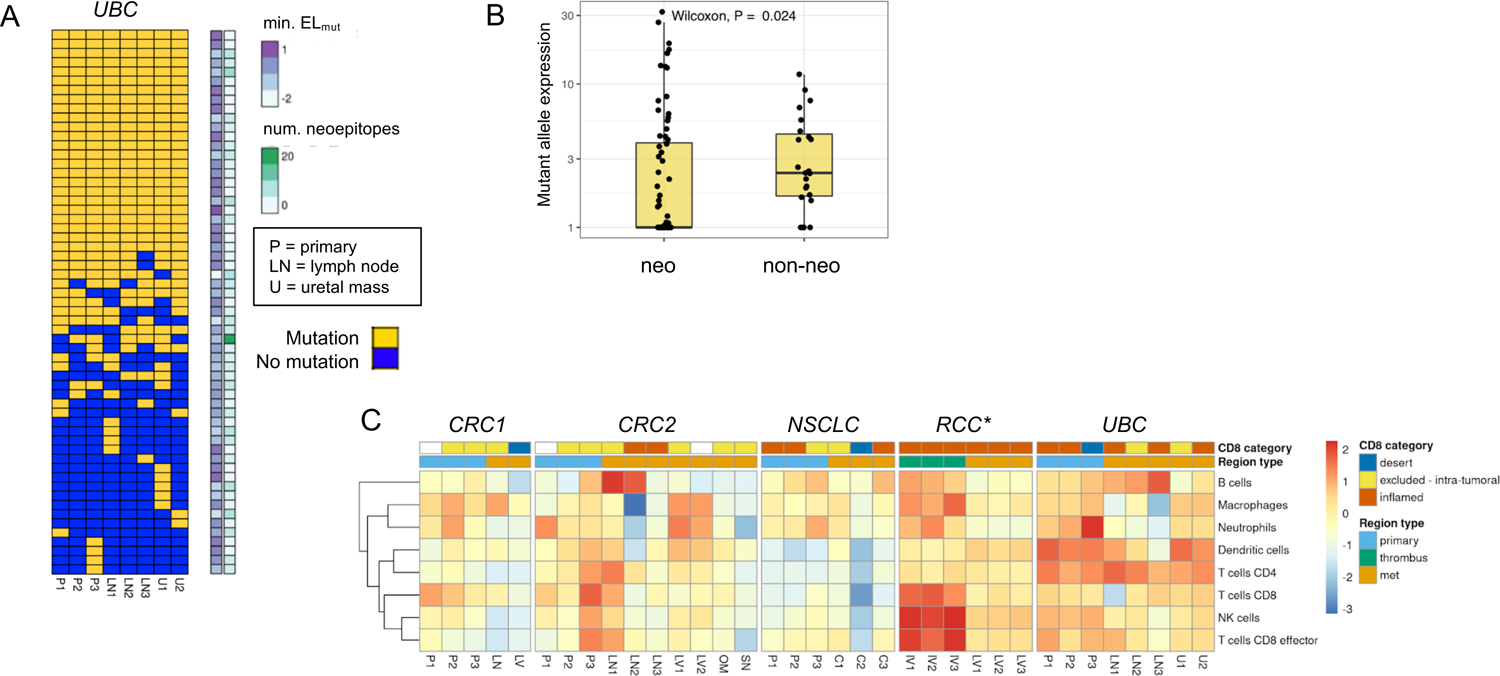
Neoantigen expression depletion and high immune inflammation across tumor regions in UBC case. (A) Sorted binary heatmap shows predicted neoantigens from the UBC case. There was no obvious relationship between neoantigen quality scores and neoantigen clonality/truncality. (B) In the UBC case, neoantigenic mutations (“neo”) had reduced mutant allele expression relative to non-neoantigenic mutations (“non-neo”). This trend was observed across all tumor regions, and was significant in 3/8 regions. (C) RNA expression signature analysis for immune cell types across tumor regions for each case. The CD8 IHC category is indicated at top, along with the type of tumor region. There was a consistent signature of inflammation across most UBC tumor samples related to dendritic cells and CD4 T-cells. *Note that RNA-seq failed for RCC primary samples, but the CD8 IHC phenotype for these regions was desert.

Notably, we did not observe similar neoantigen depletion trends in any of the other four cases (Supplementary fig. 8). We also found a consistent gene expression signature of inflammation in the UBC samples composed of dendritic cells and CD4 T cells (Fig. 6C), consistent with immune-based selection driving neoantigen loss as has been previously observed ^15^ despite heterogeneity in the CD8-based IHC classification. We looked for evidence of similar neoantigen depletion in two independent UBC cohorts, the TCGA BLCA cohort ^36^ and the IMvigor210 clinical trial cohort ^47^. In the IMvigor210 cohort, tumors with putative neoantigen expression depletion were present but relatively rare (5-8%, Supplementary fig. 9 A-C). This rarity was not surprising as most tumors may harbor only one or two strongly immunogenic neoantigens, whereas a tumor would need to have several immunogenic neoantigens undergoing expression downregulation for neoantigen depletion to be detected following this approach. The signal of neoantigen depletion was also potentially present but less apparent in the TCGA BLCA cohort (Supplementary fig. 9 D-F).

Finally, we looked across the five cases for evidence of loss of neoantigen presentation via HLA allele loss, which has been shown to be prevalent in NSCLC (McGranahan Cell 2017). Using a custom method for detecting HLA loss, we observed clonal single-allele loss of HLA-A/B/C genes in the NSCLC tumor via copy neutral LOH, and non-clonal single-allele loss of HLA-A/B/C in the liver metastasis of the RCC tumor via genomic deletion (Supplementary fig. 10). The NSCLC tumor was only modestly and somewhat heterogeneously immune-cell infiltrated (as assessed by RNA-seq and IHC) across regions (Fig. 6C), and it also did not have especially high tumor mutation burden (6.5-7.5 muts/Mb across regions). Interestingly, the immune phenotype of the RCC liver metastasis was inflamed by both IHC and RNA-seq, consistent with the metastasis-specific HLA loss occurring as a result of immune recognition specific to that lesion.

## Discussion

iNeST relies on detecting and targeting somatic cancer mutations or neoantigens, but the factors underlying effective neoantigen targeting for anti-tumor activity are still coming into focus. Here we have provided insight into how intratumoral heterogeneity and patterns of tumor evolution across indications impact neoantigen-specific therapies. Overall our study suggests that a thorough understanding of region-to-region genetic variation in tumors may be important both to maximize the efficacy of iNeST neoantigen targeting strategies and to inform biomarkers for cancer immunotherapy.

It is commonly assumed that, to ensure efficacy, iNeST should preferentially target clonal neoantigens, as they occur in all tumor cells. Our results suggest that a more nuanced “clonality strategy” may be necessary. They highlight that the prevalence of clade-specific neoantigens in certain indications, the utility of emp-CCF to enrich for globally clonal neoantigens from single tumor samples, and the possibility of reduced expression levels of immunogenic neoantigens are all important considerations. Our cross-indication phylogenetic analyses demonstrate that focusing on clonal neoantigens would likely be effective in metastatic NSCLC and UBC, as these indications have an abundance of clonal neoantigens. Owing to the high number of clonal neoantigens in melanoma, we expect this to hold true in that indication as well. However, in indications with lower neoantigen loads and in which early branching evolution is common, such as RCC and CRC, tumors tend to harbor small numbers of clonal neoantigens. The early branching evolution of these tumors can lead to relatively large numbers of clade-specific neoantigens, representing over half of all shared neoantigens in two of the tumors in our study (CRC2 and RCC). Thus, our results suggest that prioritizing high VAF mutations based on single sample sequencing in order to target clonal neoantigens would overlook many targetable clade-specific neoantigens that were not present in that sample, particularly in RCC and CRC.

The standard clonality metric, CCF, can be calculated either empirically using linear normalization of VAF to tumor purity and copy number, or with informatics tools that rely on Bayesian clustering of VAF and also generally adjust for tumor purity and local copy number. In our comparisons of these metrics, VAF and emp-CCF appeared largely equivalent, with emp-CCF having marginally better association with global mutation clonality (Fig. 4B and C). An important caveat is that CCF was not evaluable for some mutations and need for post hoc adjustments in calculating emp-CCF due to the requirement for overlap of mutations and CNA calls, and also due to potentially incorrect estimation of VAF, integer CNA values, and/or tumor purity (*CN_m_* in Eq. 4, see Methods). In most tumors, only a small percentage of mutations require VAF adjustment by local copy number. This points to a tradeoff between achieving quantification accuracy for that small fraction of mutations (which requires manual intervention and is very laborious), or potentially simplifying the process of global mutation enrichment and simply relying on VAF alone. In contrast, clust-CCF performed poorly in our comparisons with global mutation clonality (Fig. 4D and E). We suggest that the analytical process (including VAF and copy number estimation by upstream tools) and the clustering methodology of most clonality estimation tools is brittle and susceptible to frequent errors in subclone quantification or mutation assignment given noisy and variable sequencing data. Even with the enriched tumor material, as well as carefully curated sample and data quality, clust-CCF still under-performed in predicting global mutation clonality. An important distinction between emp-CCF and clust-CCF that may partly explain this poor performance is that clust-CCF values within a single sample are effectively discrete whereas emp-CCF values remain continuous. This discretization, when combined with reduced “VAF resolution” due to sample quality issues inherent to FFPE material, may lead to the poor correlation between clust-CCF and the number of regions in which a mutation is present. A potential biological confounder of this analysis is polyclonal tumor evolution, whereby metastases are seeded by multiple clones from the primary tumor leading to the presence of “shared subclonal” mutations across lesions ^11^. However, this biological phenomenon is expected to reduce the clonality-predictive value of both VAF and CCF. Therefore, either VAF alone or emp-CCF can be used to enrich clonal mutations and neoantigens, and future work is needed toward predictive metrics for globally clonal mutations that result from formally modeling tumor evolutionary processes.

Our finding that samples with fewer mutations tend to enrich for globally clonal mutations suggests that there may be a general tradeoff between sensitivity for mutation detection and enriching for globally clonal mutations. The lymph node samples in the NSCLC case illustrate how poor sample quality or insufficient tumor material can lead to the appearance of globally clonal mutation enrichment while in fact many globally clonal mutations go undetected. Similarly, samples with higher numbers of mutations may capture late-emerging subclones, which can sometimes seed metastases. On the one hand this can lead to these samples having lower proportions of global mutations, but on the other hand it could in some patients point to clade-specific neoantigens. Understanding when distinct subclones will emerge as metastases will be key.

Single tumor samples are most commonly used for biomarker exploration and therapeutic development in iNeST. Our focus on specimen quality assurance, extensive sampling and tumor enrichment afforded us a data set capable of shedding more light on the impact of regional sampling in the detection of neoantigens. We found that targeting primary tumors over metastases would give us the highest likelihood of identifying globally clonal neoantigens across indications. Sequencing a second sample would further improve clonal neoantigen detection across indications with the greatest impact in indications demonstrating increased branching. Additionally, samples with low tumor content tended to suffer from mutation under-detection due to insufficient tumor input. Lymph node metastases were regions most likely to demonstrate small tumor regions resulting in insufficient tumor content or area of harvest for mutation detection in our data set. Therefore, we show that iNeST efforts to target clonal neoantigens may benefit from sampling primary tumor regions that provide the highest level of tumor content and/or acquiring a second sample in certain indications (when clinical decision making allows it).

Our systematic characterization of neoantigen quality scores, neoantigen expression, and HLA loss in five metastatic tumors revealed distinct mechanisms with the potential to impact iNeST efficacy. We found minimal evidence for genetic loss of neoantigens via CNA in the five multi-region cases. However, given the limited number of patients examined, genetic neoantigen loss may still occur and further studies with larger sample sizes are warranted. The statistically significant neoantigen expression depletion observed in the UBC case, and the related trends in two additional UBC cohorts, suggested that in some patients certain neoantigens may show decreased expression due to immune surveillance and concomitant negative selection imposed on the tumor. The selective depletion of neoantigenic mutations and the resulting depletion across all tumor regions suggested that the neoantigen depletion in this UBC tumor may have been established early in tumor development and largely maintained into metastatic disease. Notably, only 1-5% of neoantigens are thought to be truly immunogenic ^48^, and so it may be that only specific neoantigens detected by the immune system relatively early in tumor development are subject to such expression depletion and that such immune detection is variable across patients. We identified HLA loss occurring both clonally in the NSCLC case and specific to the liver metastasis in the RCC case. Although both the IVC tumor thrombus and the liver metastasis had an immune-inflamed CD8 IHC phenotype, the liver metastasis did have the highest number of neoantigens overall, suggesting one possible reason for the observed HLA loss. Further studies are needed to explore whether presentation of specific neoantigens by tumor cells leads to neoantigen depletion subsequent to immune detection, and to determine whether such neoantigen depletion impacts tumor response to immunotherapy.

We note that depending on the level and type of neoantigen depletion or HLA loss, one might expect that a given tumor might be primed for response to immunotherapy due to the presence of expanded neoantigen-specific T-cells prior to therapy (if neoantigen presentation is reduced just below a threshold required for a robust anti-tumor response), or the tumor might be rendered refractory to immunotherapy if neoantigen presentation is reduced to a point where it cannot readily be restored. Thus, it remains unclear whether neoantigens that have been subject to immune-based depletion constitute good targets for individualized immunotherapies. However, we note that in our study and in previous studies, tumors with signs of immune-editing via neoantigen depletion or HLA allele loss tend to be immune inflamed, consistent with the possibility that these tumors could remain responsive to iNeST or checkpoint inhibitor therapy. Taken together, our results suggest the following may be necessary to ensure iNeST efficacy across indications: (1) consider the likely mode of tumor evolution per indication and the inclusion of clade-specific neoantigens in indications with branching evolution such as RCC, and (2) consider neoantigens of various expression levels, and (3) consider the presence/absence of the presenting HLA allele in the tumor when prioritizing neoantigen targets. Our efforts to characterize neoantigen qualities and presentation function anticipate a broader and deeper collective effort to understand how selective pressures sculpt the tumor neoantigen landscape and ultimately to associate hallmark patterns of neoantigen depletion with potential for response to immunotherapy treatments.

## Acknowledgments

We thank our colleagues for helpful discussions. Funding was provided by Genentech.

## Lo et al. Supplementary Figures

**Supplementary Fig. 1:**
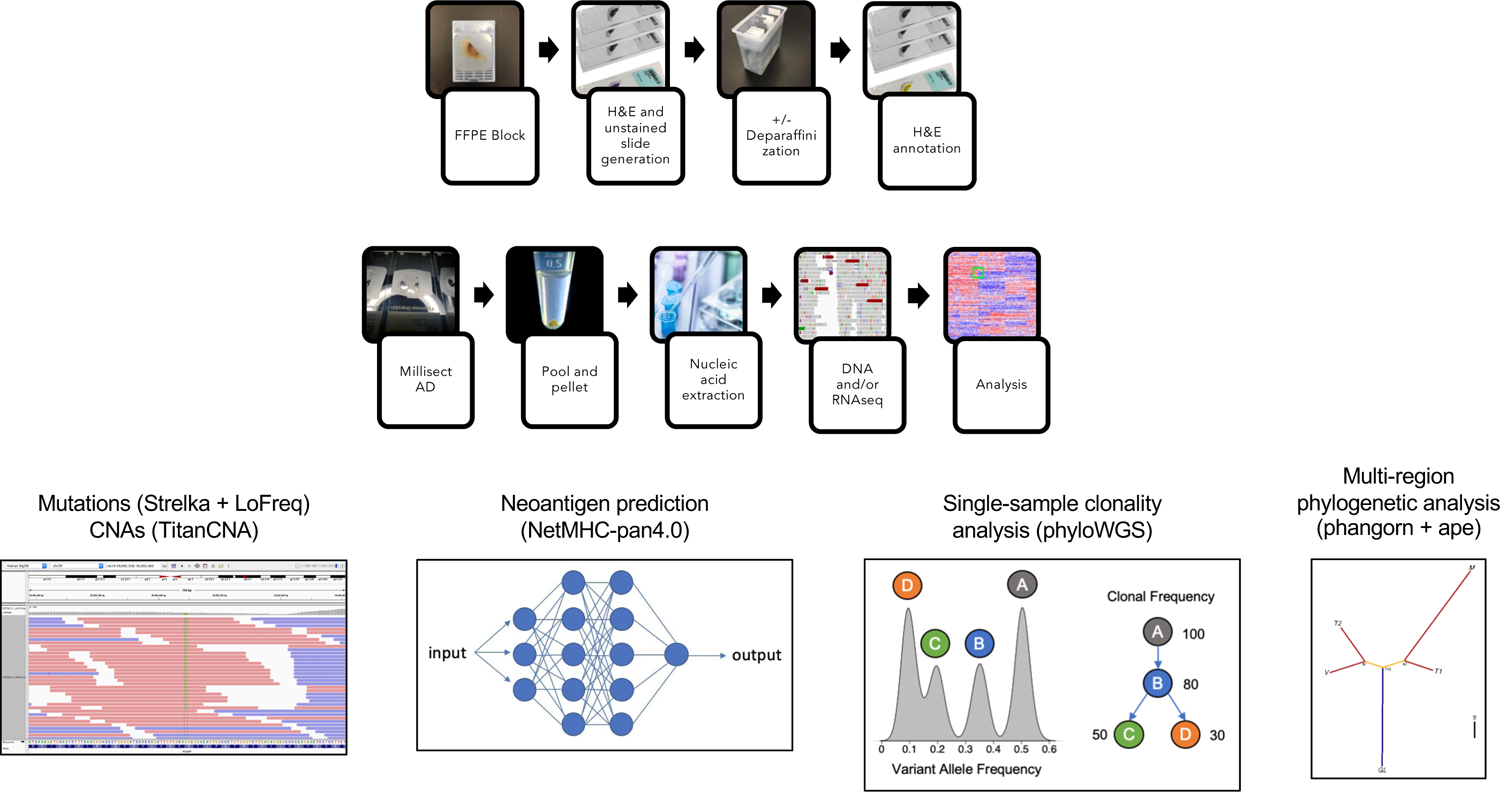
Tissue processing, MR-seq and data analysis workflow. H&E and unstained slides were generated to annotate the tumor region and perform tumor enrichment using the AVENIO Millisect prior to nucleic acid extraction, sequencing and analysis as shown to identify all somatic mutations and subsequently predict neoantigens.

**Supplementary Fig. 2:**
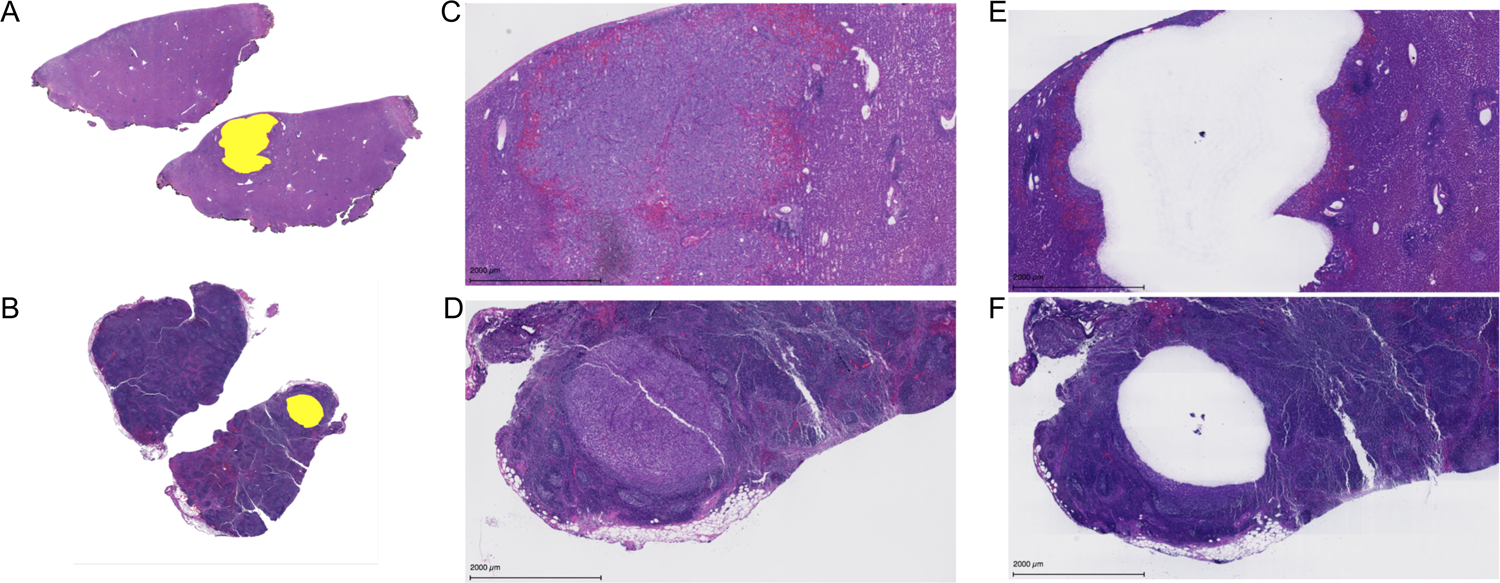
Tumor enrichment using AVENIO Millisect semi-automated tissue dissection. Annotated digital masks (A, B) were created from H&E slides to mask tumor regions (B, C) and applied to serially sectioned unstained slides for milling of selected areas for tumor enrichment. H&E stained post-dissection slides (E, F) demonstrates tumor regions that were successfully dissected for entry into nucleic acid extraction and sequencing.

**Supplementary Fig. 3:**
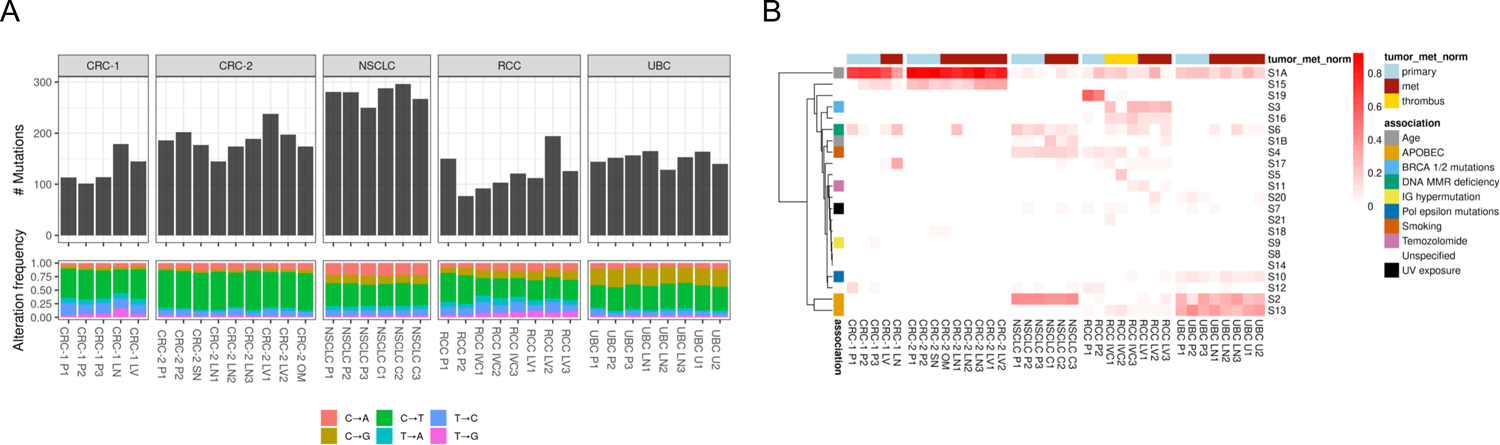
Mutation class frequencies and somatic mutation signatures across tumor regions. (A) Somatic mutation counts and SNV mutation flanking nucleotide contexts for all tumor regions of all patients. (B) Relative contribution of established somatic signatures (Alexandrov et al 2013) to the SNV mutational spectra of each region across all patients.

**Supplementary Fig. 4:**
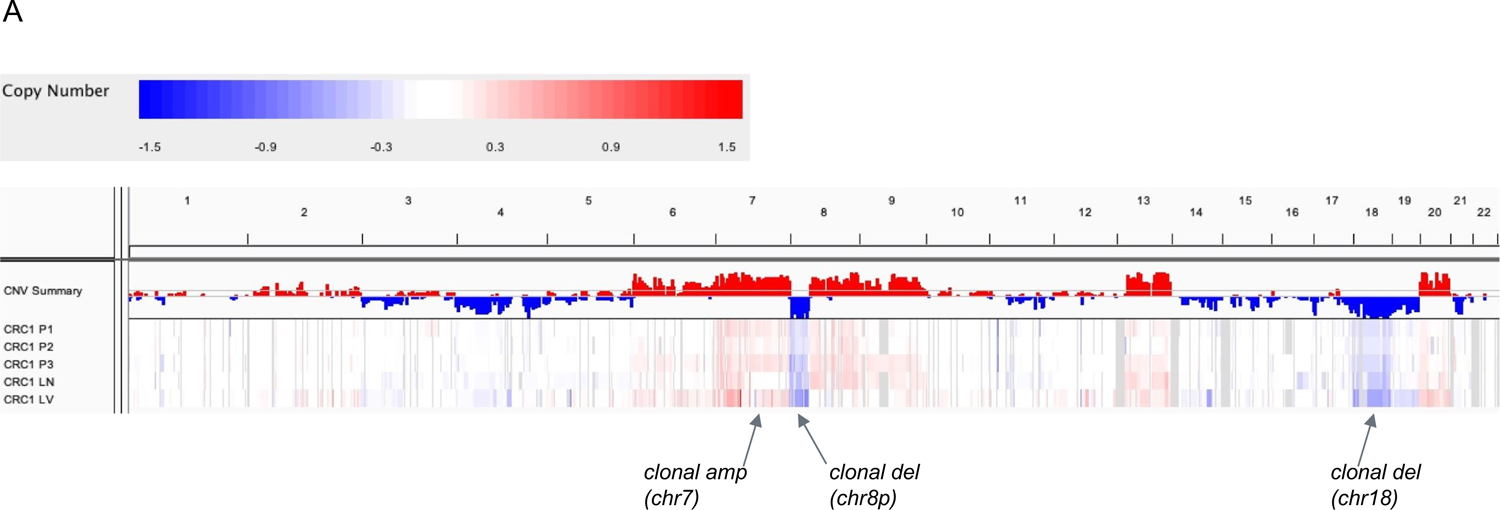
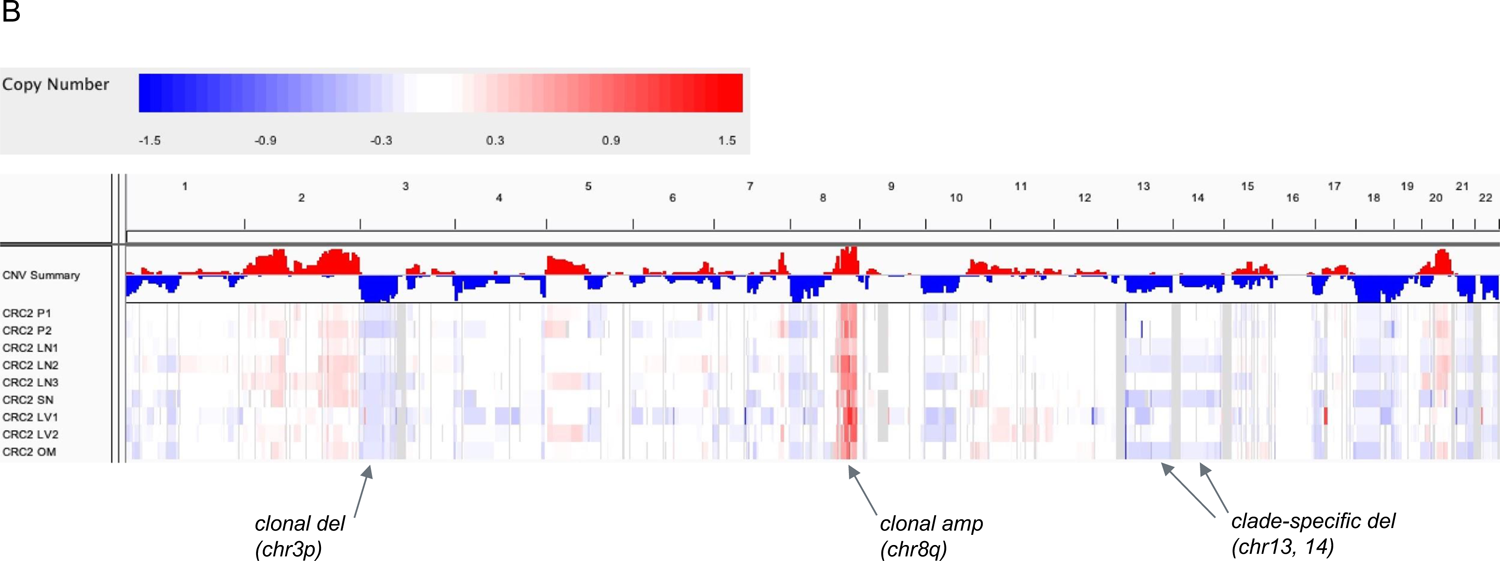
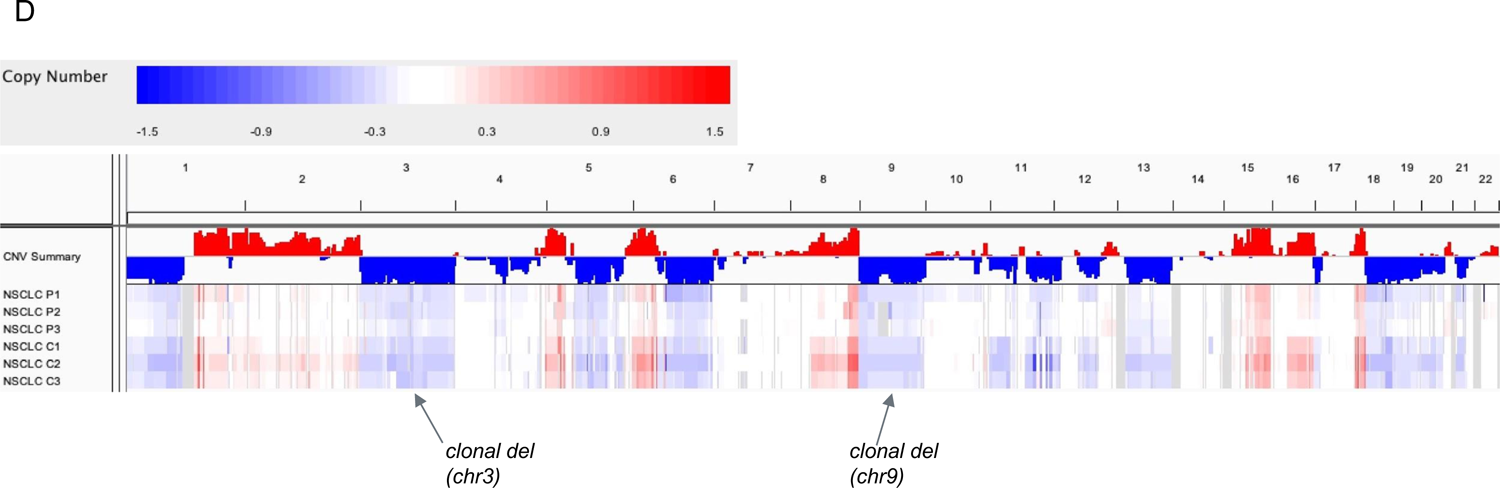
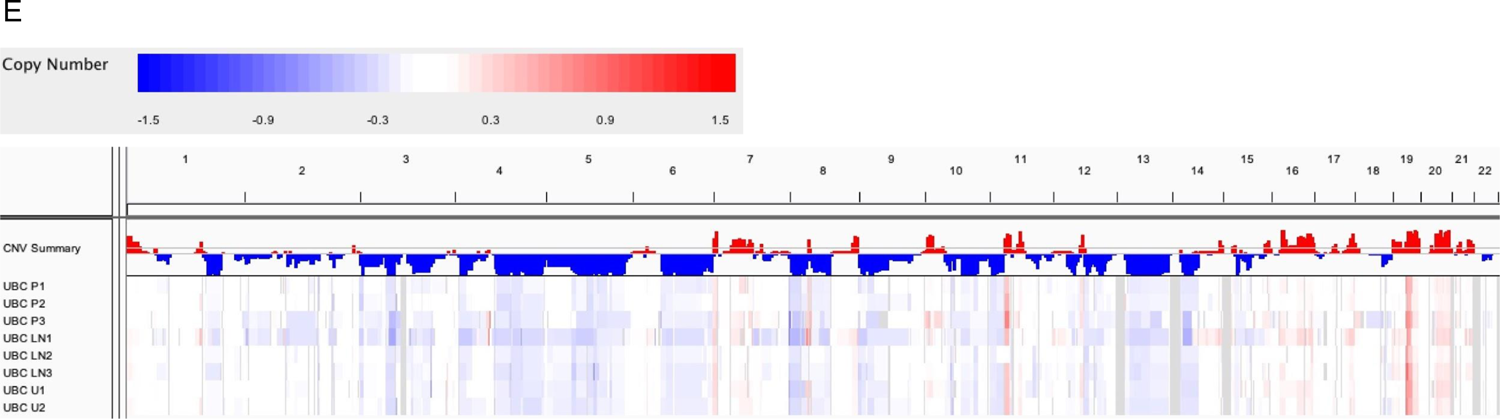
Copy number analysis of the five multi-region cases. (A) Copy number analysis of the CRC1 tumor samples. IGV plot of median logR values for copy number segments from TitanCNA. Specific CNAs with known association with CRC are indicated. (B) Copy number analysis of the CRC2 tumor samples. IGV plot of median logR values for copy number segments from TitanCNA. Specific CNAs with known association with CRC are indicated. (C) Copy number analysis of the RCC tumor samples. IGV plot of median logR values for copy number segments from TitanCNA. Specific CNAs with known association with RCC are indicated. (D) Copy number analysis of the NSCLC tumor samples. IGV plot of median logR values for copy number segments from TitanCNA. Specific CNAs with known association with NSCLC are indicated. (E) Copy number analysis of the UBC tumor samples. IGV plot of median logR values for copy number segments from TitanCNA.

**Supplementary Fig. 5:**
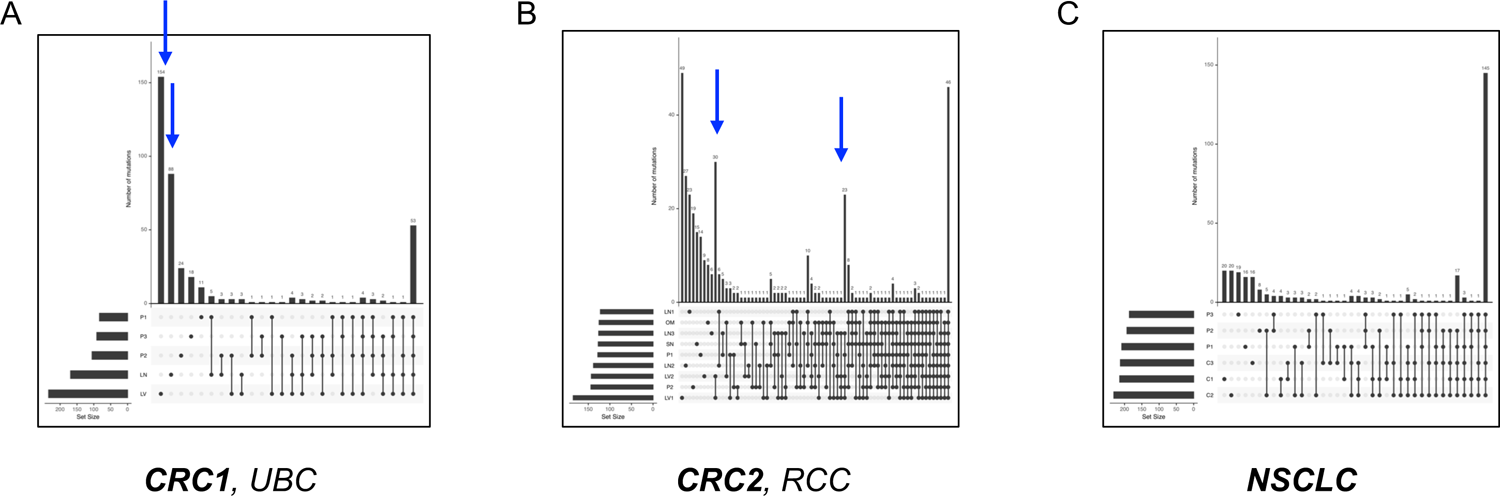
UpSetR analyses to address how and when sequencing a second biopsy would help to identify clonal mutations. A second biopsy can eliminate “singleton” mutations, enriching for clonal mutations. At the same time, sample quality is key to ensure sensitive mutation detection. (A) The CRC1 and UBC cases would benefit from two biopsies, eliminating the singleton mutations highlighted by the blue arrows (the CRC1 case is shown). (B) In the CRC2 and RCC cases, a second biopsy would enrich for clonal mutations, but cannot distinguish clade-specific from clonal mutations (the CRC2 case is shown). (C) The NSCLC would have minimal benefit from a second biopsy due to few singletons or lesion-specific mutations.

**Supplementary Fig. 6.**
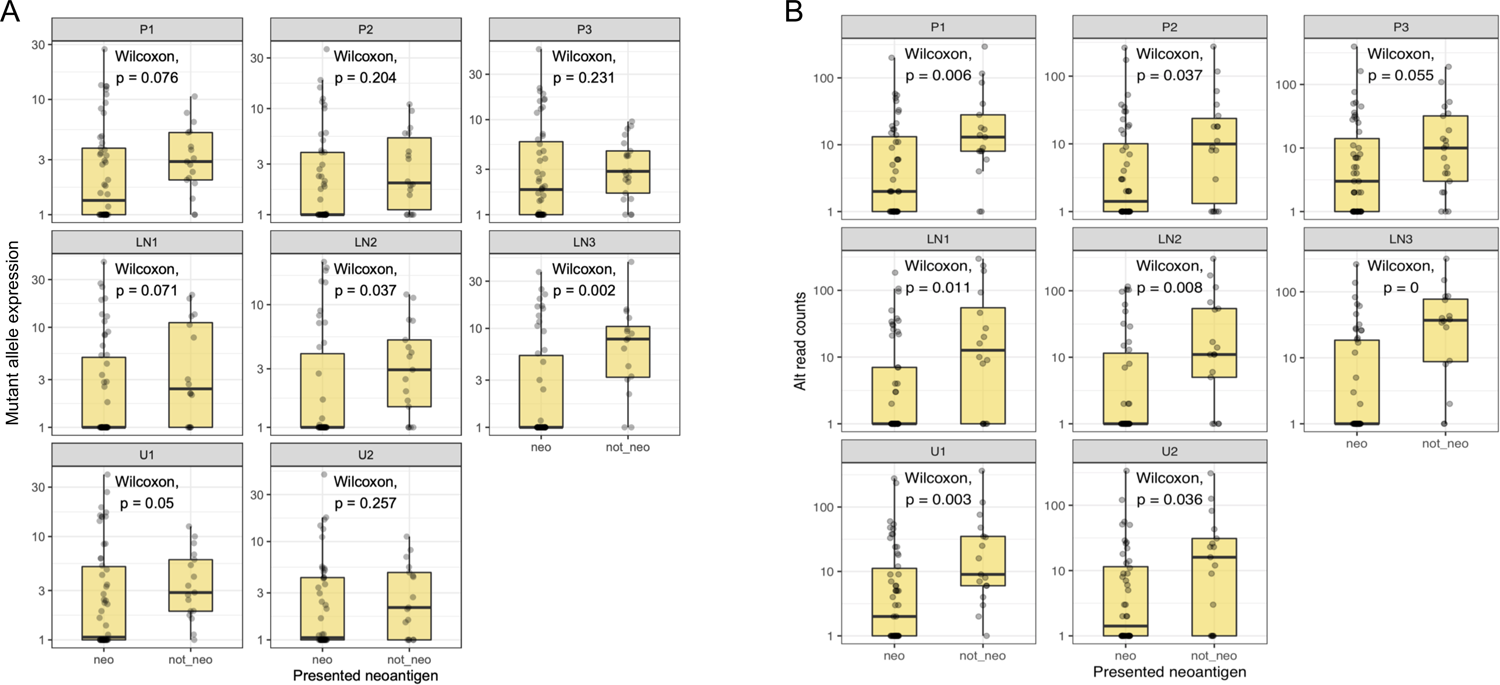
(A) Analysis of mutant allele expression versus neoantigen status for all somatic mutations across UBC tumor regions, and (B) analysis of alt-allele read counts versus neoantigen status for all somatic mutations across UBC tumor regions.

**Supplementary Fig. 7.**
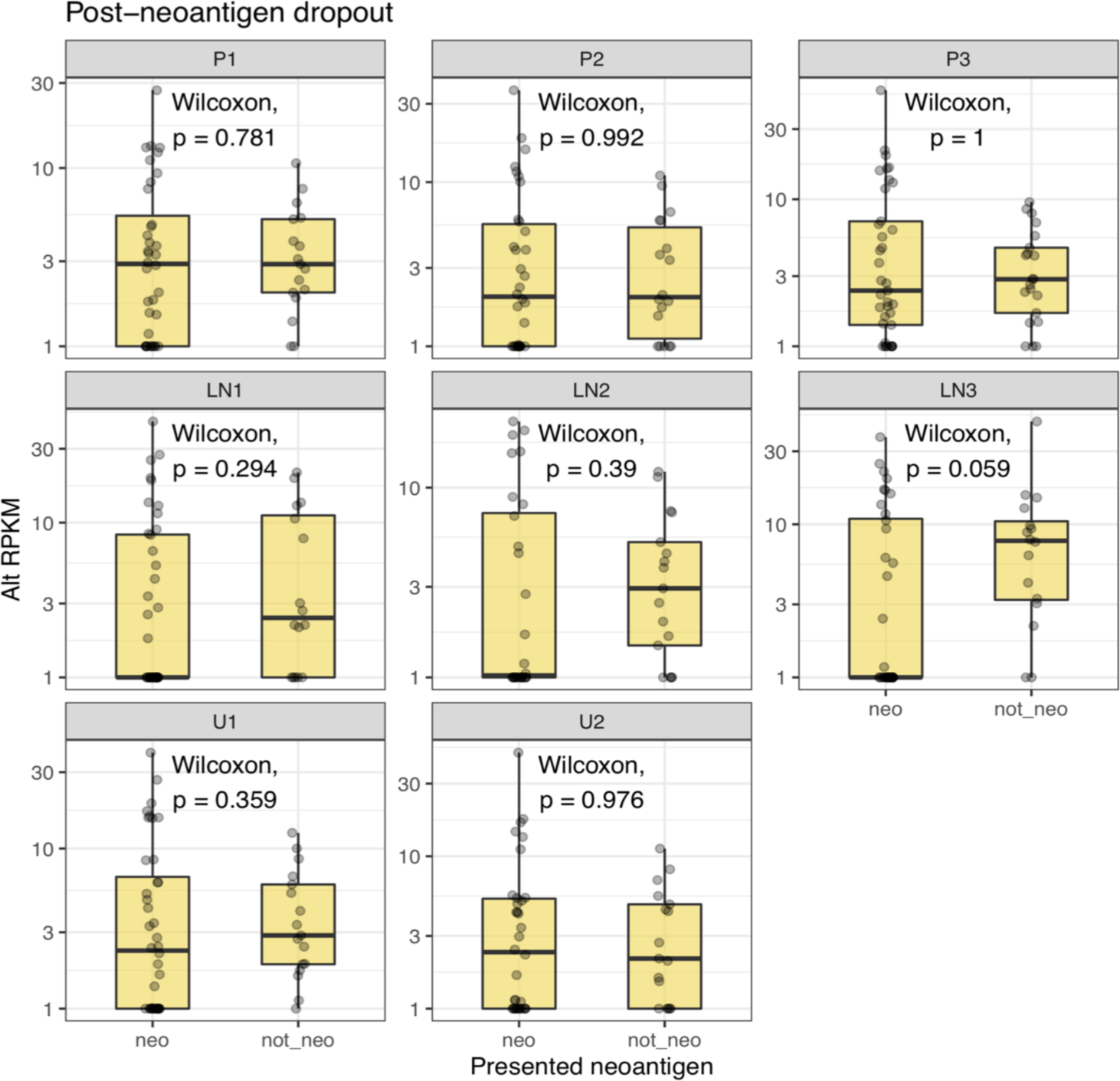
Analysis of mutant allele expression versus neoantigen status after removing the 14 candidate neoantigens mediating the expression depletion effect across UBC tumor regions.

**Supplementary Fig. 8.**
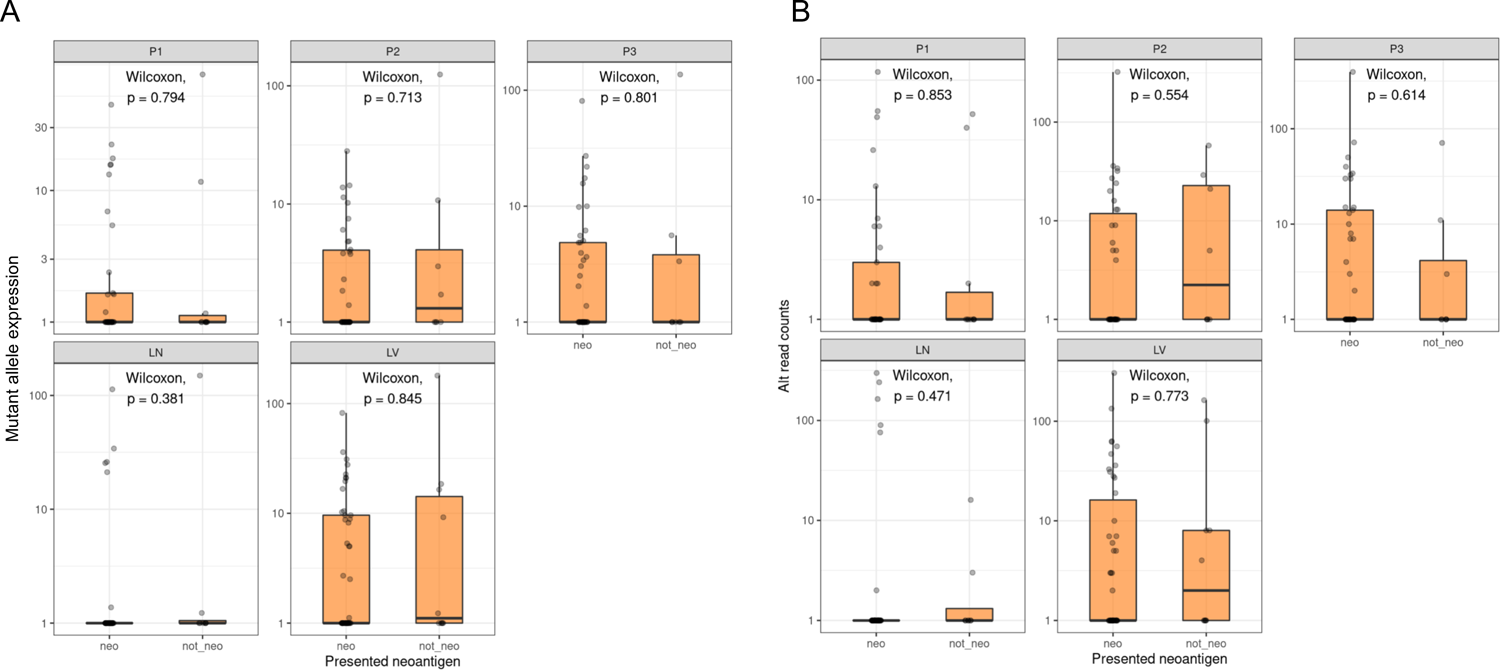
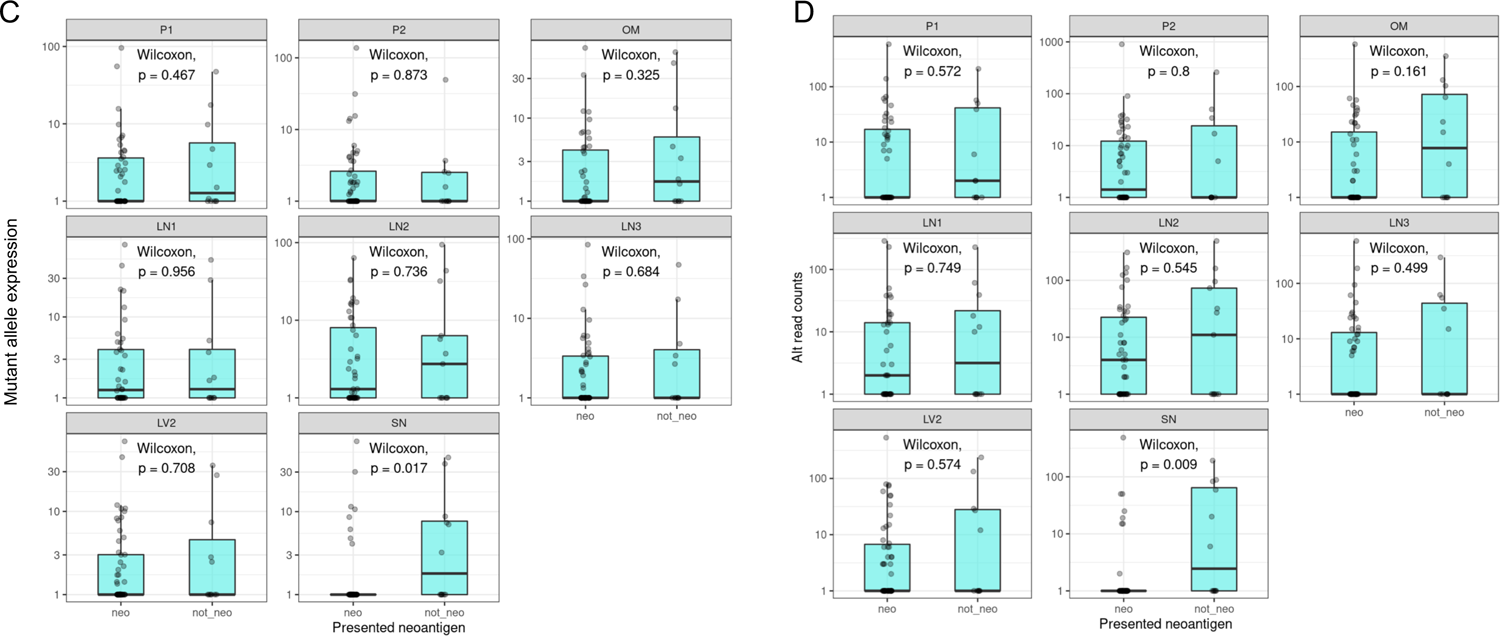
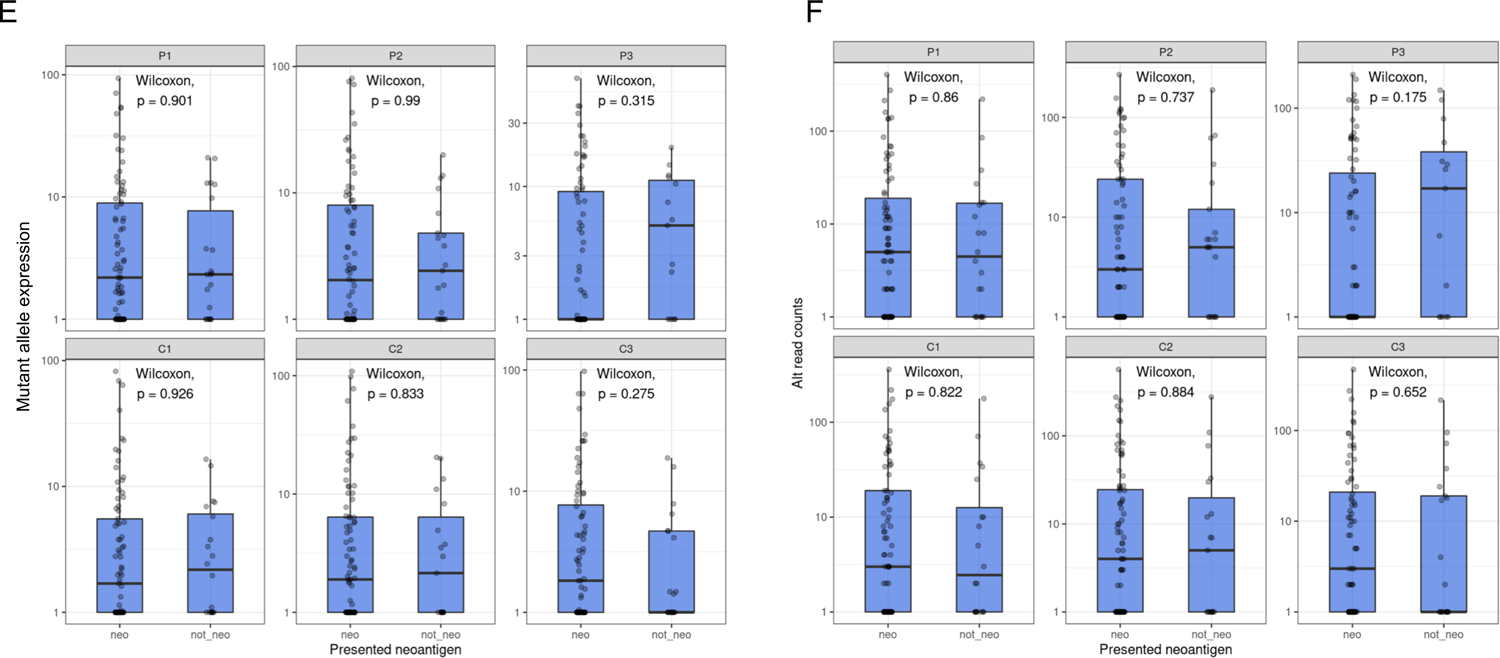
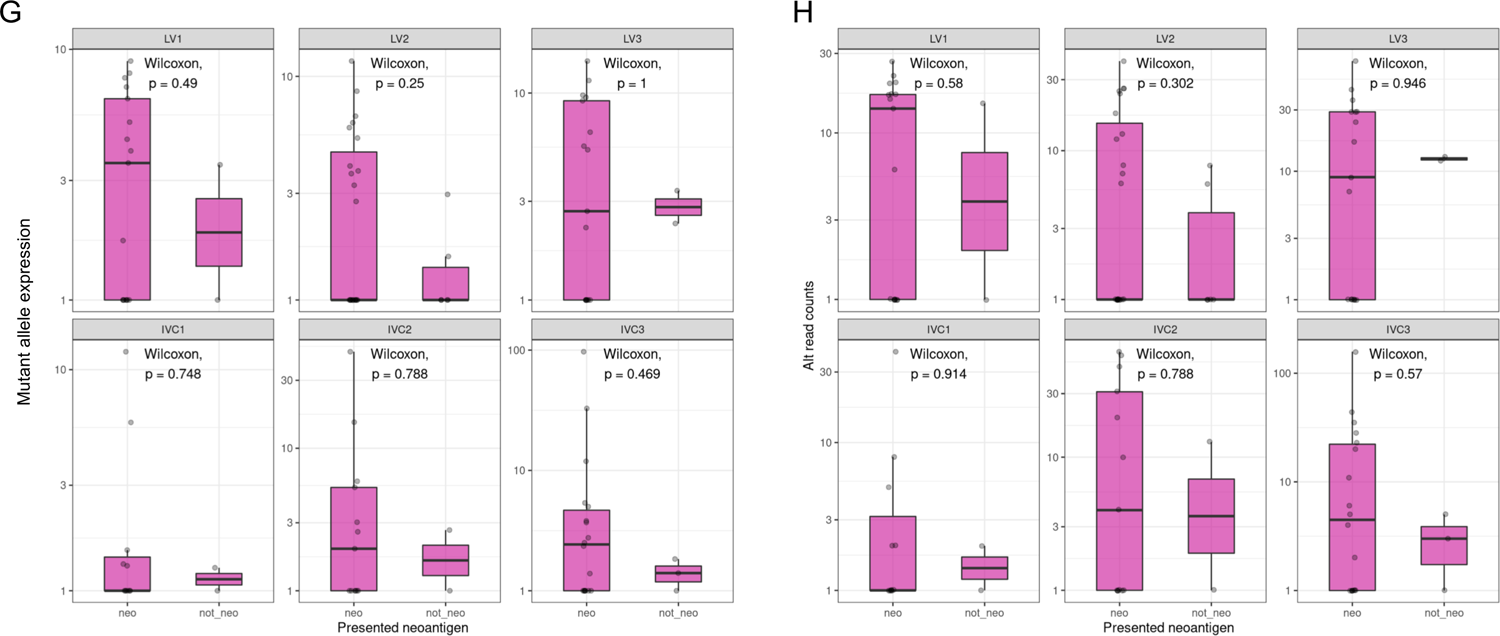
(A) Analysis of mutant allele expression versus neoantigen status for all somatic mutations across CRC1 tumor regions, and (B) analysis of alt-allele read counts versus neoantigen status for all somatic mutations across CRC1 tumor regions. (C) Analysis of mutant allele expression versus neoantigen status for all somatic mutations across CRC2 tumor regions, and (D) analysis of alt-allele read counts versus neoantigen status for all somatic mutations across CRC2 tumor regions. (E) Analysis of mutant allele expression versus neoantigen status for all somatic mutations across NSCLC tumor regions, and (F) analysis of alt-allele read counts versus neoantigen status for all somatic mutations across NSCLC tumor regions. (G) Analysis of mutant allele expression versus neoantigen status for all somatic mutations across RCC tumor regions, and (H) analysis of alt-allele read counts versus neoantigen status for all somatic mutations across RCC tumor regions.

**Supplementary Fig. 9:**
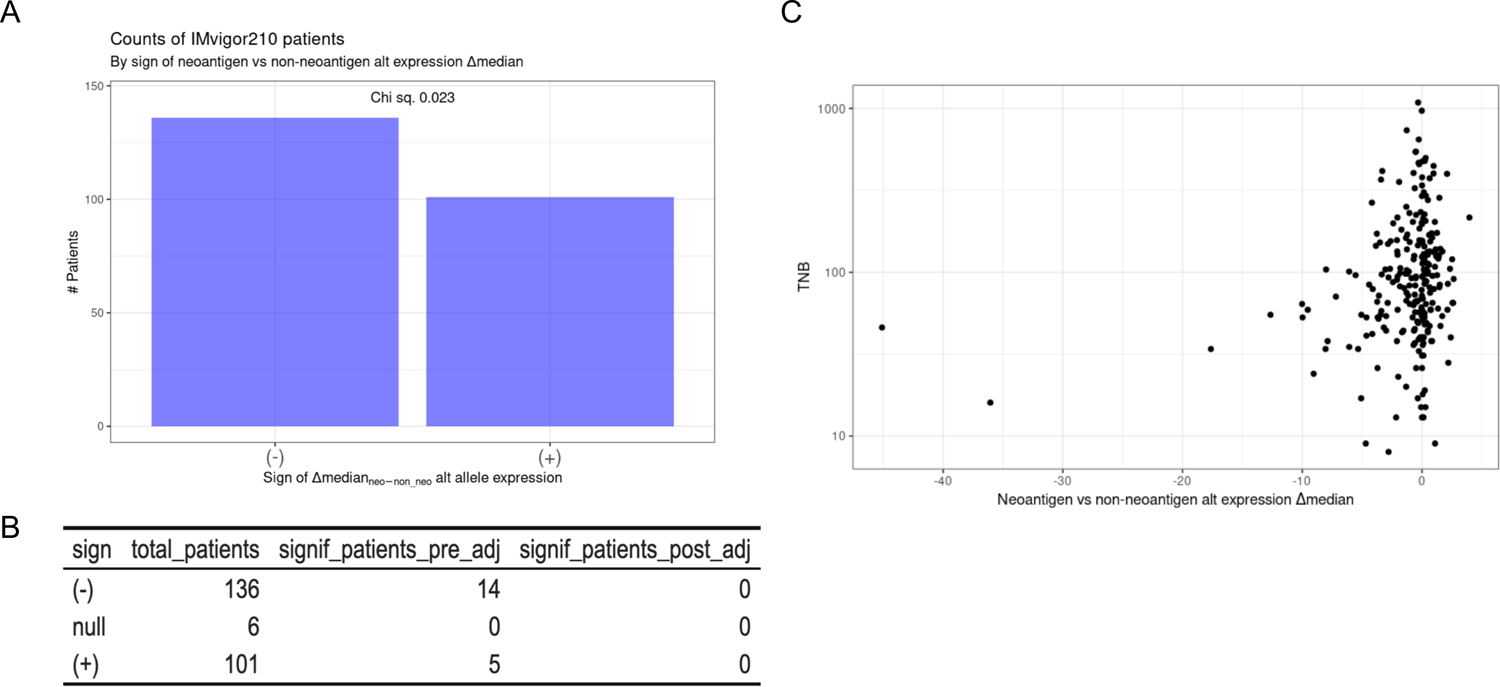
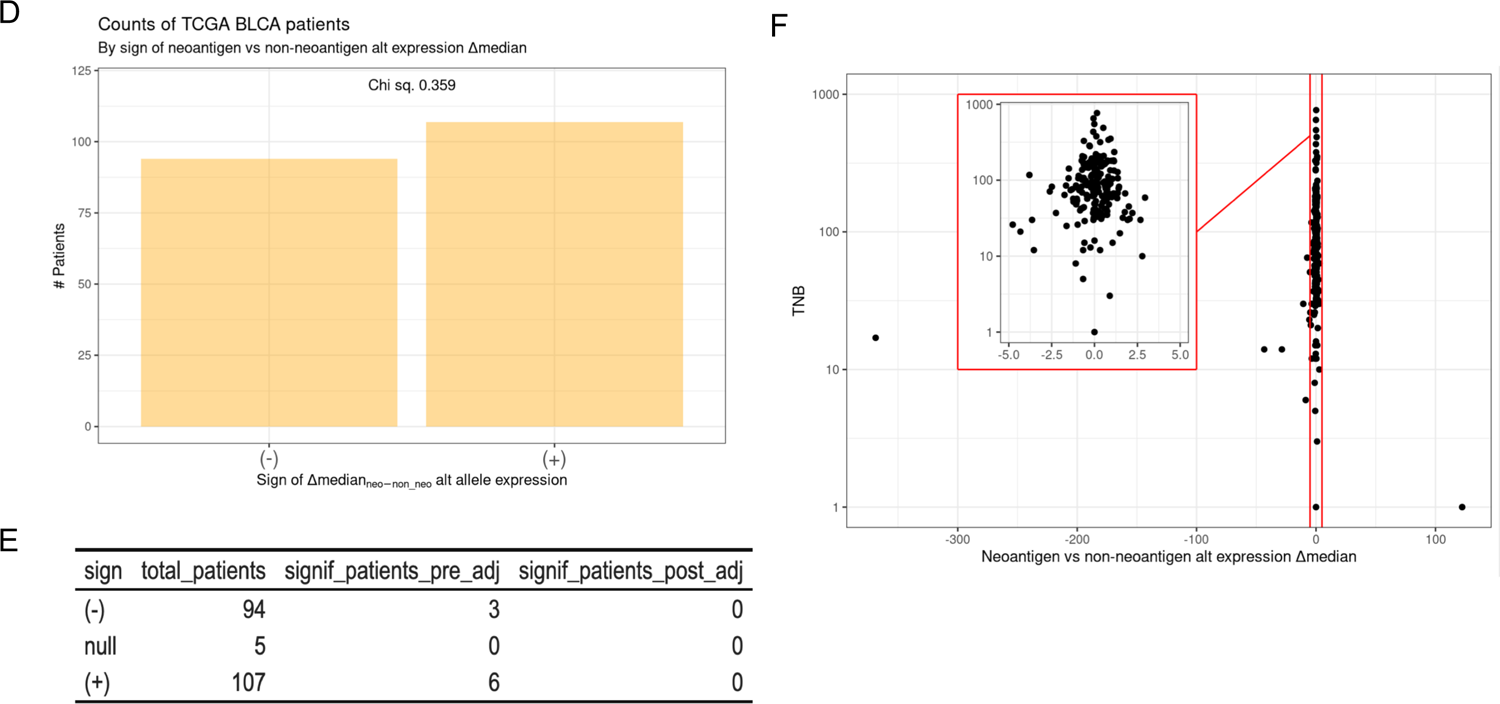
Evidence for neoantigen expression depletion in the TCGA BLCA and IMvigor210 bladder cancer cohorts. (A) The relationship between neoantigen status and mutant allele expression was examined for UBC patients in the IMvigor210 cohort (n=243). The difference in median expression for neoantigenic (“neo”) mutations versus median expression for non-neoantigenic (“non-neo”) mutations was determined for each patient. Barplot shows the number of patients with a negative sign or a positive sign from this Lmedian analysis, with a Chi-square p-value from a test of the proportions. There were 11 patients with only neo or non-neo mutations and these were excluded from the analysis. (B) The table shows the specific numbers of patients with each Lmedian sign along with the numbers of patients that individually showed significant differences in their “neo” versus “non-neo” expression levels (by a Wilcoxon rank-sum test). (C) Scatter plot of tumor neoantigen burden (“TNB”) versus Lmedian for each patient. Note the leftward trend of Lmedian, indicative of samples with neoantigen expression depletion. (D) and (E) The relationship between neoantigen status and mutant allele expression was examined for UBC patients in the TCGA BLCA cohort (n=202). Similar analyses are shown as in (A) and (B).

**Supplementary Fig. 10:**
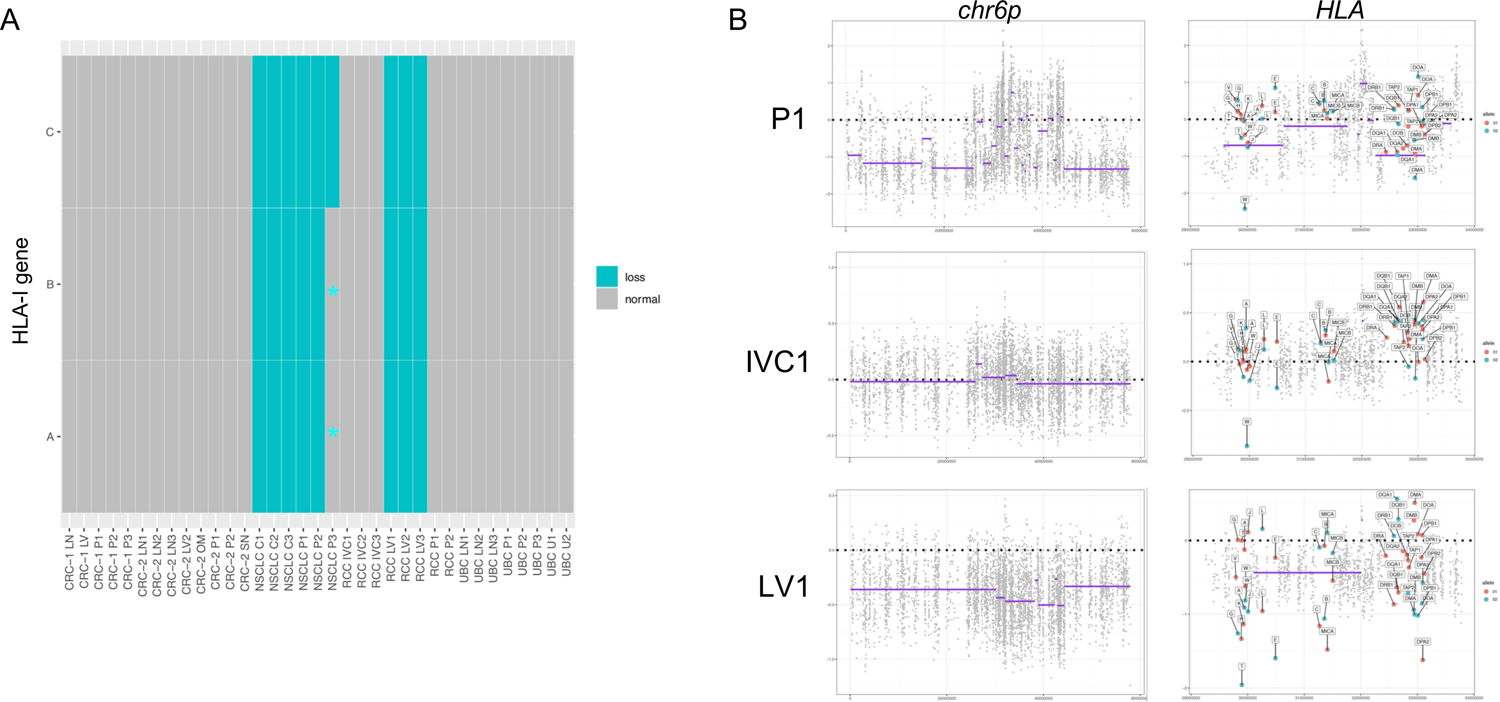
Allele-specific copy number alterations in HLA-I genes across all tumor regions. (A) Single allele HLA loss was detected for the HLA-A/B/C genes in all NSCLC samples, and in the liver metastasis samples from the RCC case. (*) indicates there was a signal of HLA loss that fell just below the significance threshold. (B) Tumor/normal log ratios across chromosome 6p and the HLA region. Purple lines indicate segment means for the non-HLA exons as determined by the copynumber::pcf() function in R. Labeled points at right indicate the allele-specific tumor/normal log ratios for both alleles of all HLA-I and HLA-II genes. Note that the HLA-A/B/C loss event occurs only in the liver metastasis sample (LV1), and not in the primary (P1) or IVC tumor thrombus (IVC1) sample.

**Supplementary Table 1.**
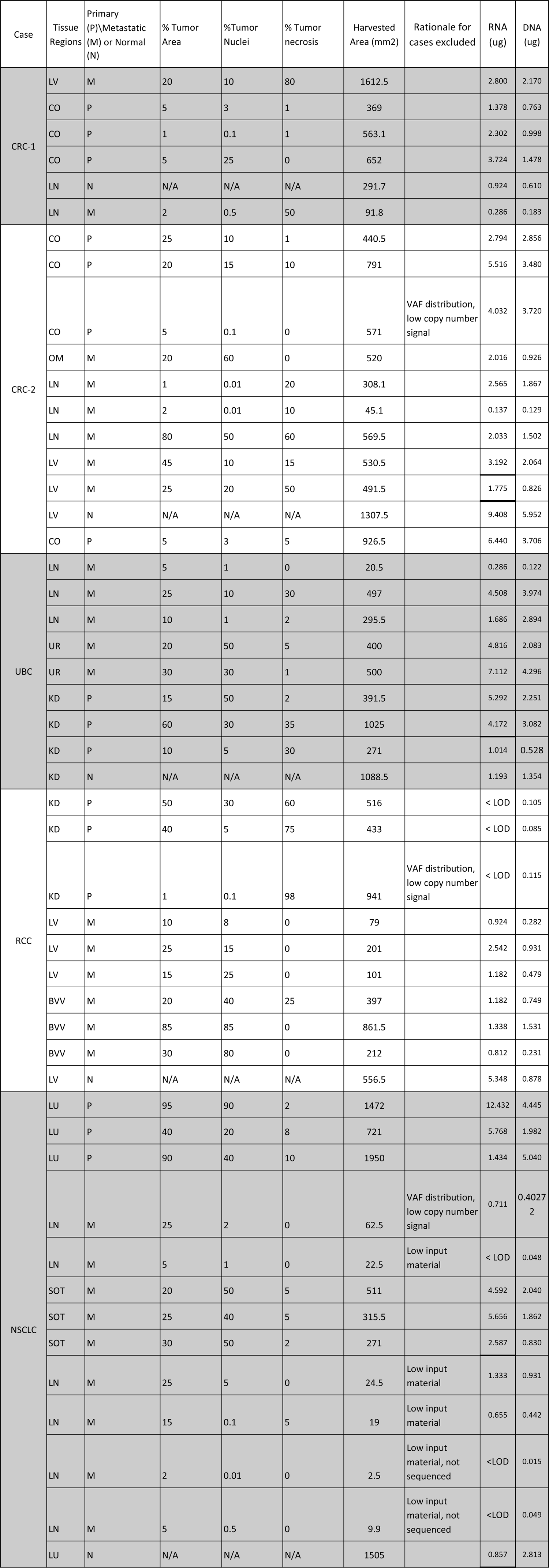
Region types, captured metrics and rationale for region exclusion, when relevant.

**Supplementary Table 2.** Numbers of total versus global mutations, expressed mutations, and neoantigens in the five multi-region cases.

|  | All mutations |  |  | Nonsynonymous mutations (missense or frameshift) |  |  | Expressed mutations |  |  | Expressed nonsynonymous mutations |  |  | Neoantigenic mutations |  |  | Expressed neoantigenic mutations |  |  |
| --- | --- | --- | --- | --- | --- | --- | --- | --- | --- | --- | --- | --- | --- | --- | --- | --- | --- | --- |
| *patient* | *total* | *global* | *global fraction* | *total* | *global* | *global fraction* | *total* | *global* | *global fraction* | *total* | *global* | *global fraction* | *total* | *global* | *global fraction* | *total* | *global* | *global fraction* |
| CRC1 | 262 | 74 | 0.28 | 131 | 44 | 0.34 | 57 | 34 | 0.60 | 35 | 26 | 0.74 | 107 | 39 | 0.36 | 31 | 23 | 0.74 |
| CRC2 | 425 | 78 | 0.18 | 201 | 36 | 0.18 | 113 | 39 | 0.35 | 63 | 20 | 0.32 | 160 | 28 | 0.18 | 51 | 15 | 0.29 |
| NSCLC | 397 | 201 | 0.51 | 166 | 101 | 0.61 | 153 | 100 | 0.65 | 84 | 59 | 0.70 | 134 | 84 | 0.63 | 66 | 49 | 0.74 |
| RCC | 375 | 31 | 0.08 | 153 | 16 | 0.10 | 89 | 19 | 0.21 | 52 | 10 | 0.19 | 129 | 15 | 0.12 | 46 | 9 | 0.20 |
| UBC | 259 | 88 | 0.34 | 117 | 50 | 0.43 | 111 | 62 | 0.56 | 60 | 35 | 0.58 | 91 | 34 | 0.37 | 42 | 22 | 0.52 |
| *total* | *1718* | *472* | *0.27* | *768* | *247* | *0.32* | *523* | *254* | *0.49* | *294* | *150* | *0.51* | *621* | *200* | *0.32* | *236* | *118* | *0.50* |

***total global fraction*** (per tumor)

**Supplementary Table 3:**
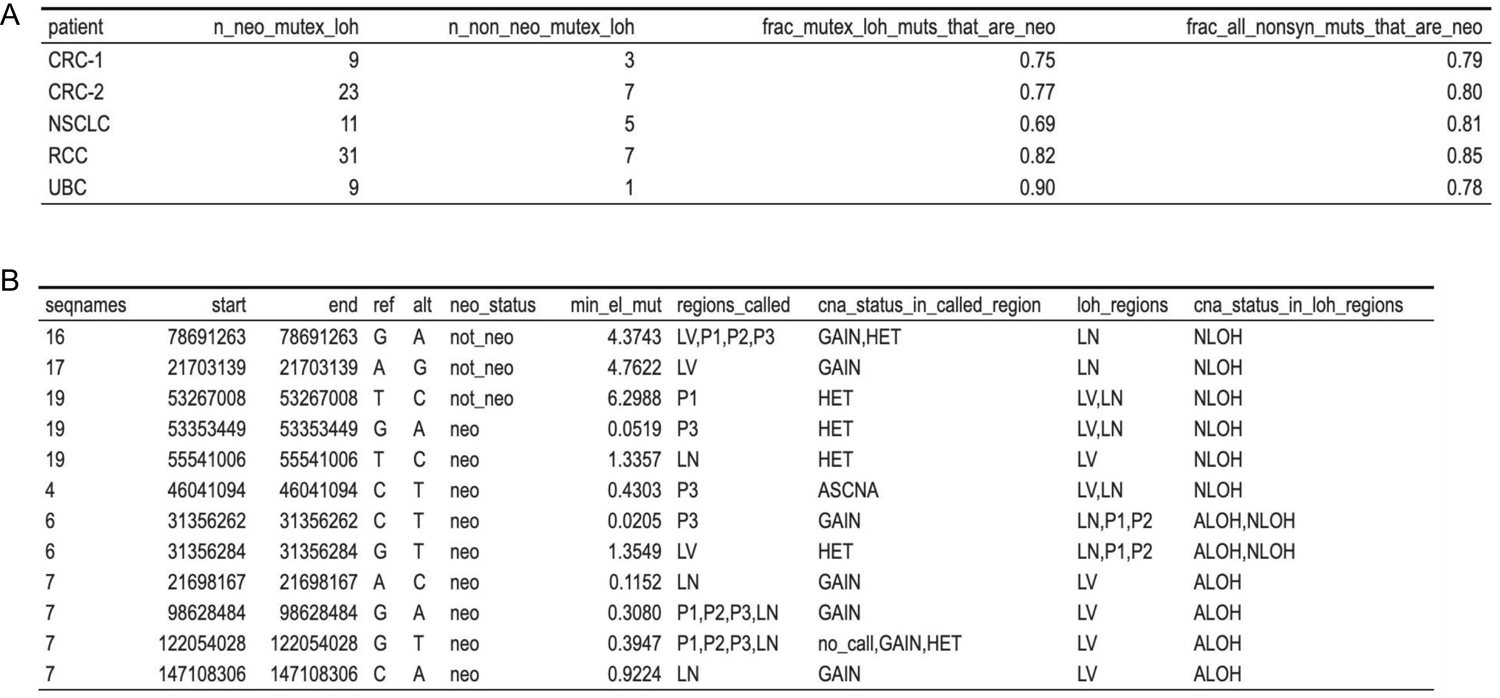
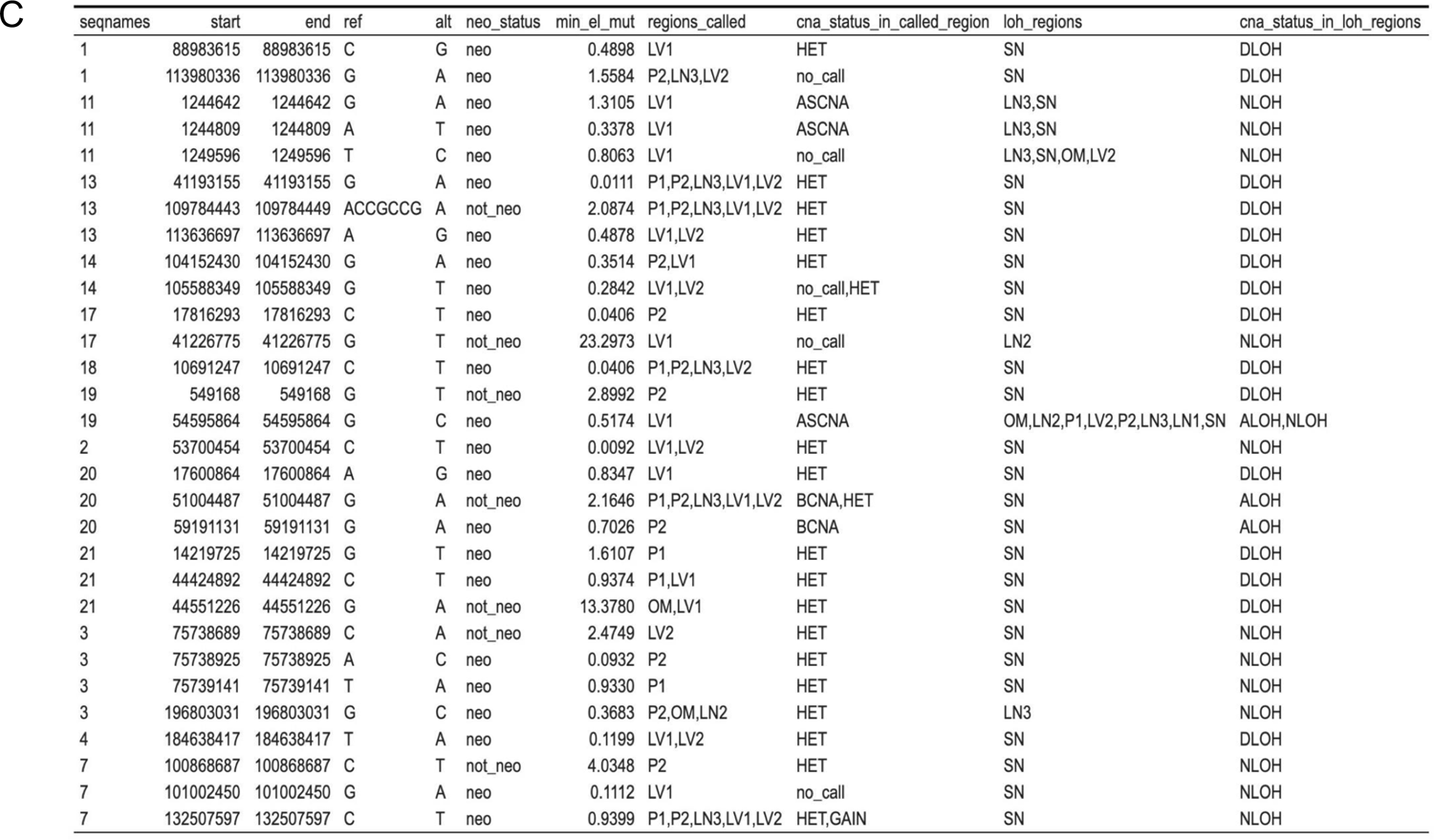
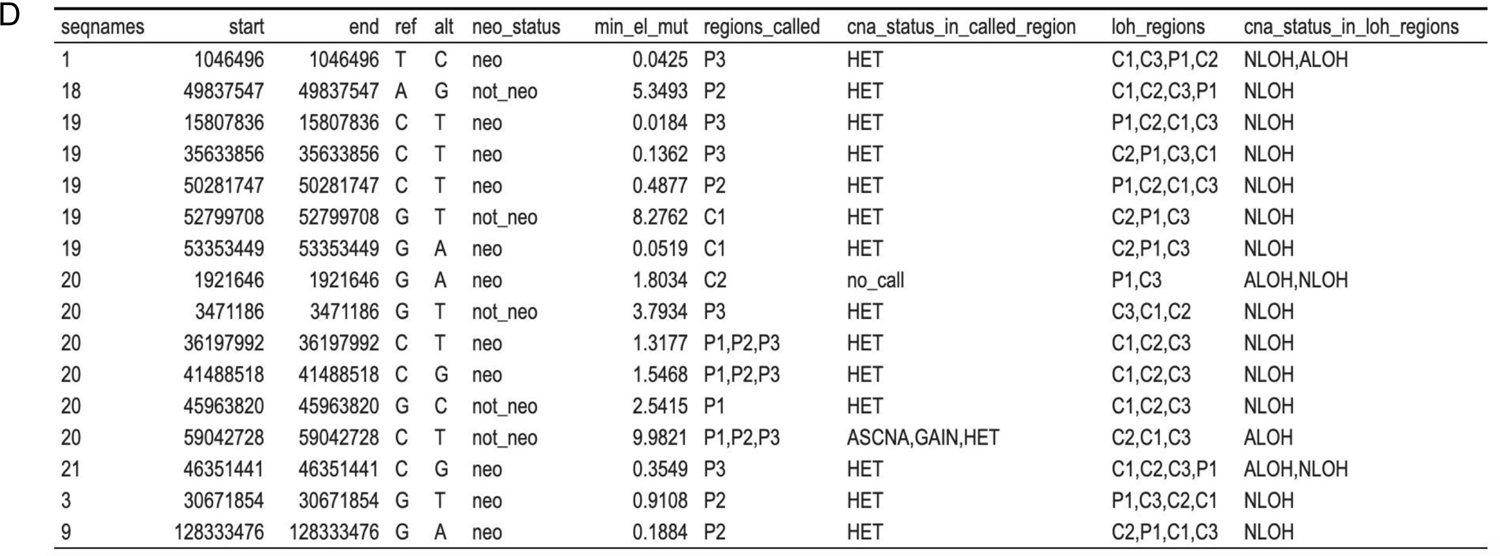
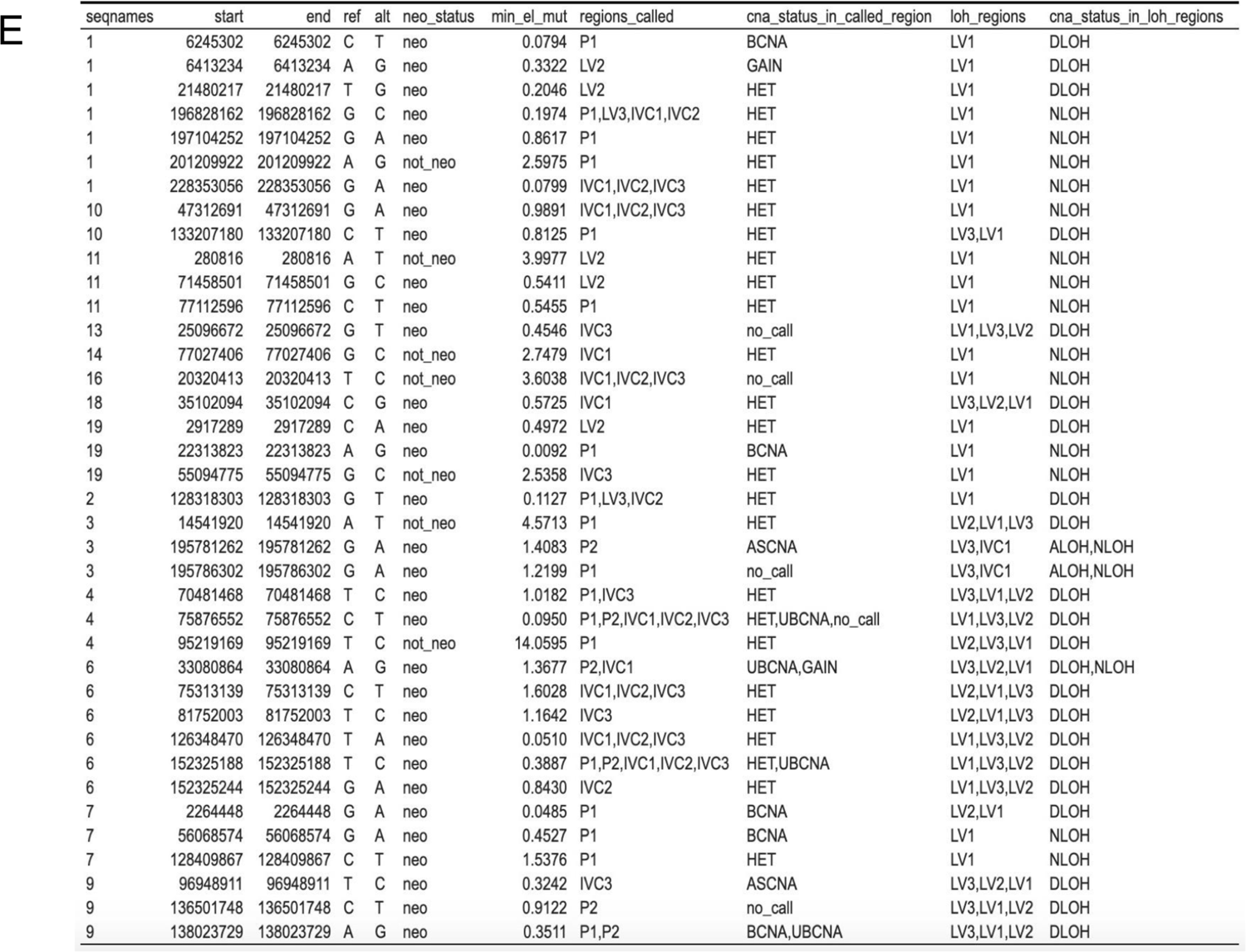
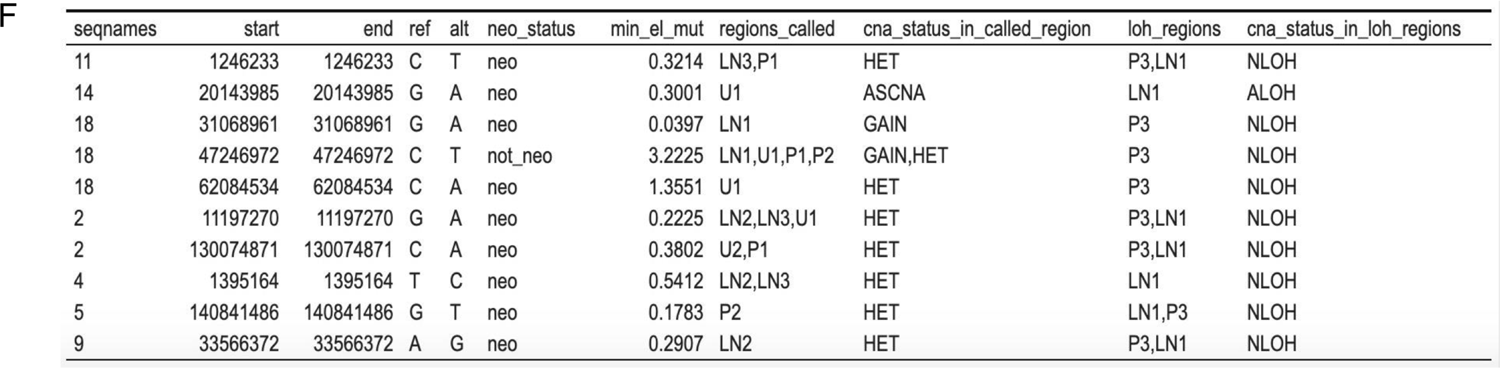
CNA based neoantigen loss in the five multi-region cases, without regard for RNAseq support of mutations. (A) Summary table showing numbers of mutations lost via genomic deletion or LOH events in each case, along with the proportion of them that are putatively neoantigenic. As a comparator, the proportion of all mutations that are putatively neoantigenic is included. (B-F) Enumeration of mutations that exhibited mutual exclusivity with LOH events in individual cases - CRC-1, CRC-2, UBC, RCC, and NSCLC respectively.

**Supplementary Table 4.** Candidate neoantigens mediating the expression depletion effect across UBC tumor regions.

| chrom | pos | ref | alt | gene | mut_peptide | min_el_mut | n_regions | expr_regions |
| --- | --- | --- | --- | --- | --- | --- | --- | --- |
| 2 | 162396927 | C | T | KCNH7 | IVVAILGKNDIFEEMVHLYAKPGKS | 0.0208 | 8 | LN1, LN2, LN3, U1, U2, P1, P2, P3 |
| 2 | 178718119 | G | A | TTN | QCKVDGTPEIRIFWYKEHTKLRSAP | 0.1764 | 8 | LN1, LN2, LN3, U1, U2, P1, P2, P3 |
| 13 | 87676004 | G | T | SLITRK5 | NNLFRFVPLTHLYLRGNRLKLLPYV | 0.2527 | 8 | LN1, LN2, LN3, U1, U2, P1, P2, P3 |
| 6 | 31871375 | C | T | SLC44A4 | QQGISGLIDSLNTRDISVKIFEDFA | 0.5174 | 8 | LN1, LN2, LN3, U1, U2, P1, P2, P3 |
| 17 | 29969201 | G | A | EFCAB5 | MLTQVEKKKVLTKADTPSKFDPINY | 0.8164 | 8 | LN1, LN2, LN3, U1, U2, P1, P2, P3 |
| 12 | 8059594 | C | G | C3AR1 | DFYGDPLENRS LQNIVQPPGEMNDR | 0.92 | 8 | LN1, LN2, LN3, U1, U2, P1, P2, P3 |
| 6 | 39879403 | G | A | DAAM2 | LPLPQDPYPSSDIPLRKKRVPQPSH | 0.9732 | 8 | LN1, LN2, LN3, U1, U2, P1, P2, P3 |
| 5 | 159084705 | T | A | EBF1 | QAIVYEGQDKNPVMCRVLLTHEIMC | 1.2098 | 8 | LN1, LN2, LN3, U1, U2, P1, P2, P3 |
| 19 | 13877273 | G | T | NANOS3 | MGTFDLWTYYLGLAHLVRALS | 0.0115 | 7 | LN1, LN2, LN3, U2, P1, P2, P3 |
| 1 | 111414895 | T | C | OVGP1 | LTPVGHQSVTPVGHQSVSPGGTTMT | 1.6476 | 7 | LN1, LN2, U1, U2, P1, P2, P3 |
| 5 | 180792305 | G | A | MGAT1 | DLEVAPDFFEYFWATYPLLKADPSL | 0.0884 | 6 | LN1, LN3, U1, P1, P2, P3 |
| 11 | 116772934 | C | G | BUD13 | MAAAPPLSKAQYLKRYLSGADAG | 0.752 | 6 | LN2, LN3, U1, U2, P1, P2 |
| 19 | 48462126 | C | A | KCNJ14 | PPPAPCFSHVASLLAAFLFALETQT | 0.0148 | 5 | LN2, LN3, U2, P1, P3 |
| 17 | 7673334 | C | G | TP53 | KPLDGEYFTLQDHTSFQKENC | 0.1465 | 5 | LN1, LN2, LN3, P1, P2 |

**Supplementary Table 5.**
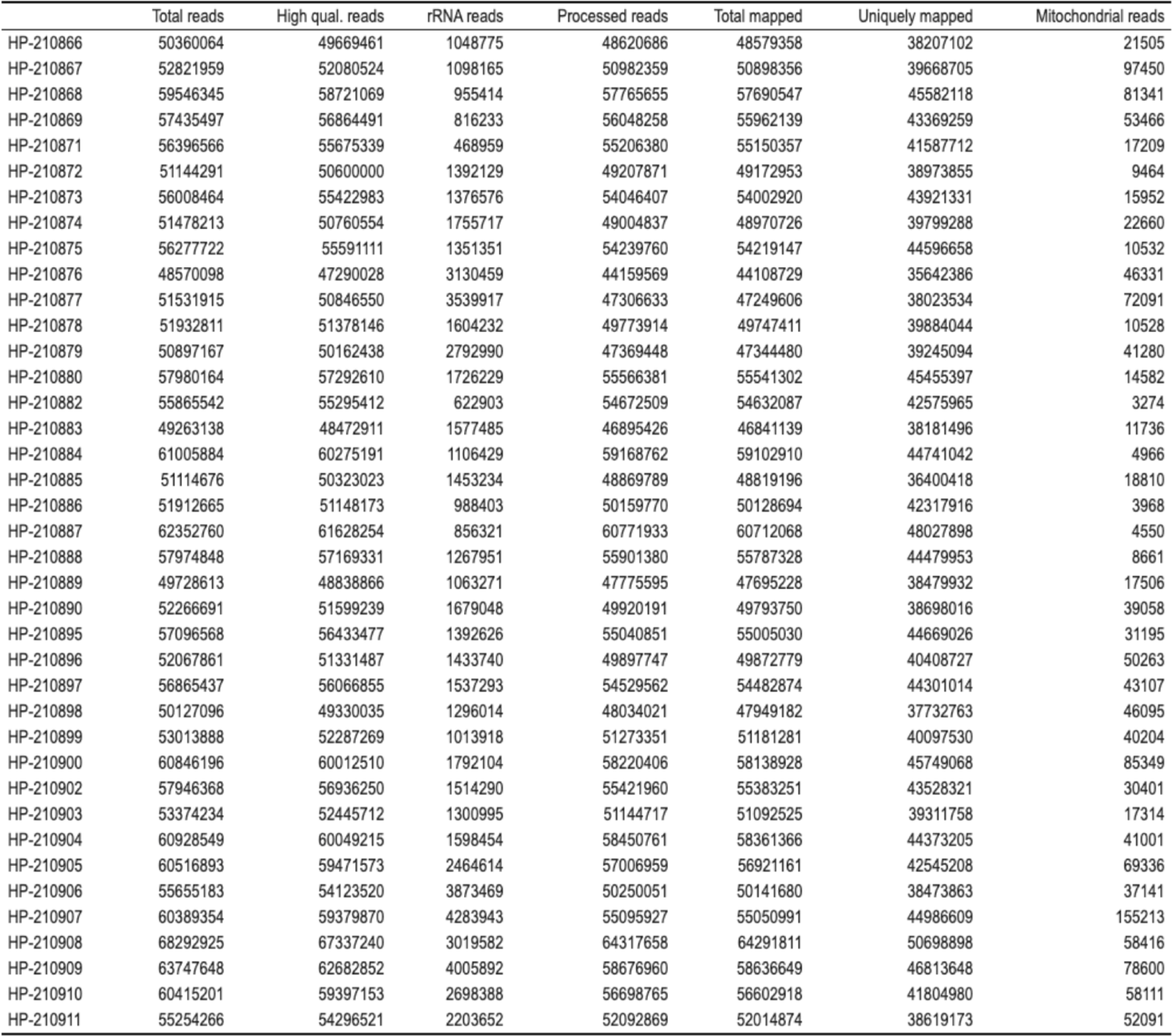
RNA-seq alignment statistics.

|  | Total reads | High qual. reads | rRNA reads | Processed reads | Total mapped | Uniquely mapped | Mitochondrial reads |
| --- | --- | --- | --- | --- | --- | --- | --- |
| HP-210866 | 50360064 | 49669461 | 1048775 | 48620686 | 48579358 | 38207102 | 21505 |
| HP-210867 | 52821959 | 52080524 | 1098165 | 50982359 | 50898356 | 39668705 | 97450 |
| HP-210868 | 59546345 | 58721069 | 955414 | 57765655 | 57690547 | 45582118 | 81341 |
| HP-210869 | 57435497 | 56864491 | 816233 | 56048258 | 55962139 | 43369259 | 53466 |
| HP-210871 | 56396566 | 55675339 | 468959 | 55206380 | 55150357 | 41587712 | 17209 |
| HP-210872 | 51144291 | 50600000 | 1392129 | 49207871 | 49172953 | 38973855 | 9464 |
| HP-210873 | 56008464 | 55422983 | 1376576 | 54046407 | 54002920 | 43921331 | 15952 |
| HP-210874 | 51478213 | 50760554 | 1755717 | 49004837 | 48970726 | 39799288 | 22660 |
| HP-210875 | 56277722 | 55591111 | 1351351 | 54239760 | 54219147 | 44596658 | 10532 |
| HP-210876 | 48570098 | 47290028 | 3130459 | 44159569 | 44108729 | 35642386 | 46331 |
| HP-210877 | 51531915 | 50846550 | 3539917 | 47306633 | 47249606 | 38023534 | 72091 |
| HP-210878 | 51932811 | 51378146 | 1604232 | 49773914 | 49747411 | 39884044 | 10528 |
| HP-210879 | 50897167 | 50162438 | 2792990 | 47369448 | 47344480 | 39245094 | 41280 |
| HP-210880 | 57980164 | 57292610 | 1726229 | 55566381 | 55541302 | 45455397 | 14582 |
| HP-210882 | 55865542 | 55295412 | 622903 | 54672509 | 54632087 | 42575965 | 3274 |
| HP-210883 | 49263138 | 48472911 | 1577485 | 46895426 | 46841139 | 38181496 | 11736 |
| HP-210884 | 61005884 | 60275191 | 1106429 | 59168762 | 59102910 | 44741042 | 4966 |
| HP-210885 | 51114676 | 50323023 | 1453234 | 48869789 | 48819196 | 36400418 | 18810 |
| HP-210886 | 51912665 | 51148173 | 988403 | 50159770 | 50128694 | 42317916 | 3968 |
| HP-210887 | 62352760 | 61628254 | 856321 | 60771933 | 60712068 | 48027898 | 4550 |
| HP-210888 | 57974848 | 57169331 | 1267951 | 55901380 | 55787328 | 44479953 | 8661 |
| HP-210889 | 49728613 | 48838866 | 1063271 | 47775595 | 47695228 | 38479932 | 17506 |
| HP-210890 | 52266691 | 51599239 | 1679048 | 49920191 | 49793750 | 38698016 | 39058 |
| HP-210895 | 57096568 | 56433477 | 1392626 | 55040851 | 55005030 | 44669026 | 31195 |
| HP-210896 | 52067861 | 51331487 | 1433740 | 49897747 | 49872779 | 40408727 | 50263 |
| HP-210897 | 56865437 | 56066855 | 1537293 | 54529562 | 54482874 | 44301014 | 43107 |
| HP-210898 | 50127096 | 49330035 | 1296014 | 48034021 | 47949182 | 37732763 | 46095 |
| HP-210899 | 53013888 | 52287269 | 1013918 | 51273351 | 51181281 | 40097530 | 40204 |
| HP-210900 | 60846196 | 60012510 | 1792104 | 58220406 | 58138928 | 45749068 | 85349 |
| HP-210902 | 57946368 | 56936250 | 1514290 | 55421960 | 55383251 | 43528321 | 30401 |
| HP-210903 | 53374234 | 52445712 | 1300995 | 51144717 | 51092525 | 39311758 | 17314 |
| HP-210904 | 60928549 | 60049215 | 1598454 | 58450761 | 58361366 | 44373205 | 41001 |
| HP-210905 | 60516893 | 59471573 | 2464614 | 57006959 | 56921161 | 42545208 | 69336 |
| HP-210906 | 55655183 | 54123520 | 3873469 | 50250051 | 50141680 | 38473863 | 37141 |
| HP-210907 | 60389354 | 59379870 | 4283943 | 55095927 | 55050991 | 44986609 | 155213 |
| HP-210908 | 68292925 | 67337240 | 3019582 | 64317658 | 64291811 | 50698898 | 58416 |
| HP-210909 | 63747648 | 62682852 | 4005892 | 58676960 | 58636649 | 46813648 | 78600 |
| HP-210910 | 60415201 | 59397153 | 2698388 | 56698765 | 56602918 | 41804980 | 58111 |
| HP-210911 | 55254266 | 54296521 | 2203652 | 52092869 | 52014874 | 38619173 | 52091 |

**Supplementary Table 6.**
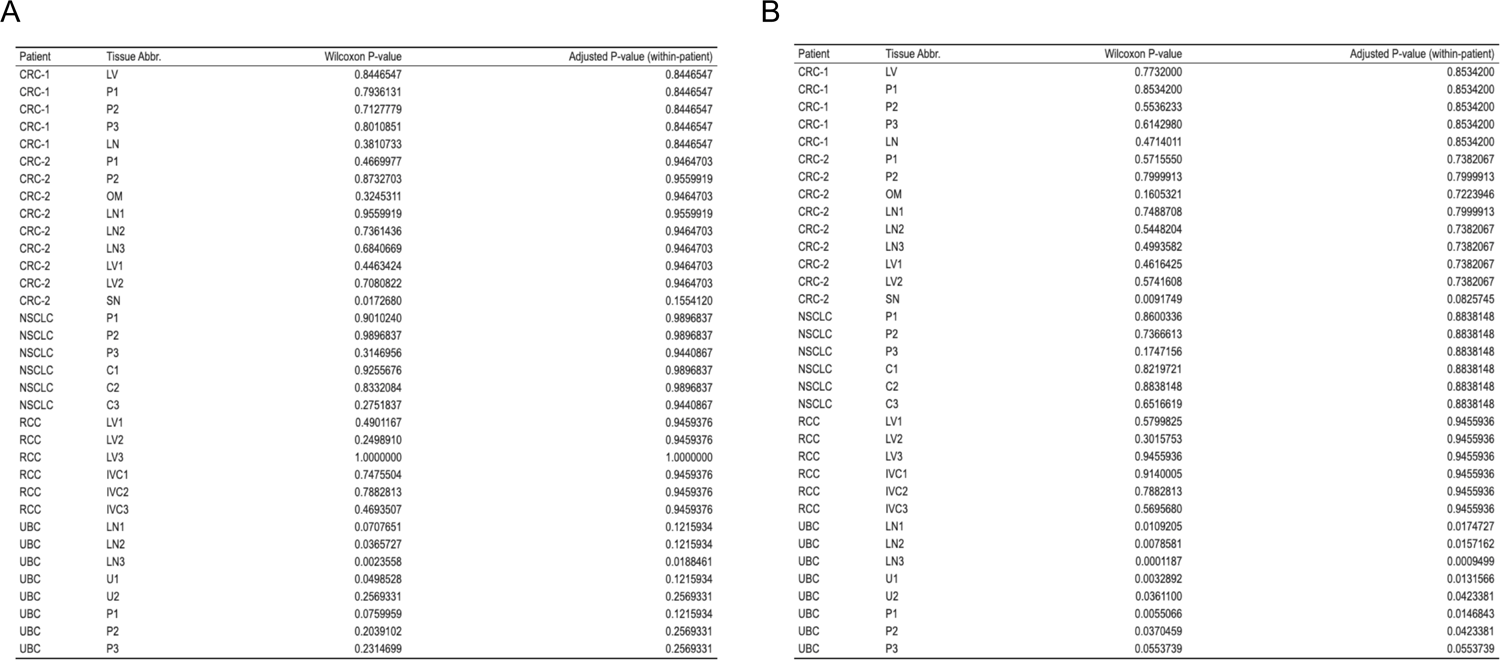
P-values and within-patient adjusted P-values for region-level neoantigen vs non-neoantigen expression comparisons using A) alt RPKM or B) alt-allele read counts.

Supplementary Tables 6A,B A
| Patient | Tissue Abbr. | Wilcoxon P-value | Adjusted P-value (within-patient) |
| --- | --- | --- | --- |
| CRC-1 | LV | 0.8446547 | 0.8446547 |
| CRC-1 | P1 | 0.7936131 | 0.8446547 |
| CRC-1 | P2 | 0.7127779 | 0.8446547 |
| CRC-1 | P3 | 0.8010851 | 0.8446547 |
| CRC-1 | LN | 0.3810733 | 0.8446547 |
| CRC-2 | P1 | 0.4669977 | 0.9464703 |
| CRC-2 | P2 | 0.8732703 | 0.9559919 |
| CRC-2 | OM | 0.3245311 | 0.9464703 |
| CRC-2 | LN1 | 0.9559919 | 0.9559919 |
| CRC-2 | LN2 | 0.7361436 | 0.9464703 |
| CRC-2 | LN3 | 0.6840669 | 0.9464703 |
| CRC-2 | LV1 | 0.4463424 | 0.9464703 |
| CRC-2 | LV2 | 0.7080822 | 0.9464703 |
| CRC-2 | SN | 0.0172680 | 0.1554120 |
| NSCLC | P1 | 0.9010240 | 0.9896837 |
| NSCLC | P2 | 0.9896837 | 0.9896837 |
| NSCLC | P3 | 0.3146956 | 0.9440867 |
| NSCLC | C1 | 0.9255676 | 0.9896837 |
| NSCLC | C2 | 0.8332084 | 0.9896837 |
| NSCLC | C3 | 0.2751837 | 0.9440867 |
| RCC | LV1 | 0.4901167 | 0.9459376 |
| RCC | LV2 | 0.2498910 | 0.9459376 |
| RCC | LV3 | 1.0000000 | 1.0000000 |
| RCC | IVC1 | 0.7475504 | 0.9459376 |
| RCC | IVC2 | 0.7882813 | 0.9459376 |
| RCC | IVC3 | 0.4693507 | 0.9459376 |
| UBC | LN1 | 0.0707651 | 0.1215934 |
| UBC | LN2 | 0.0365727 | 0.1215934 |
| UBC | LN3 | 0.0023558 | 0.0188461 |
| UBC | U1 | 0.0498528 | 0.1215934 |
| UBC | U2 | 0.2569331 | 0.2569331 |
| UBC | P1 | 0.0759959 | 0.1215934 |
| UBC | P2 | 0.2039102 | 0.2569331 |
| UBC | P3 | 0.2314699 | 0.2569331 |

B
| Patient | Tissue Abbr. | Wilcoxon P-value | Adjusted P-value (within-patient) |
| --- | --- | --- | --- |
| CRC-1 | LV | 0.7732000 | 0.8534200 |
| CRC-1 | P1 | 0.8534200 | 0.8534200 |
| CRC-1 | P2 | 0.5536233 | 0.8534200 |
| CRC-1 | P3 | 0.6142980 | 0.8534200 |
| CRC-1 | LN | 0.4714011 | 0.8534200 |
| CRC-2 | P1 | 0.5715550 | 0.7382067 |
| CRC-2 | P2 | 0.7999913 | 0.7999913 |
| CRC-2 | OM | 0.1605321 | 0.7223946 |
| CRC-2 | LN1 | 0.7488708 | 0.7999913 |
| CRC-2 | LN2 | 0.5448204 | 0.7382067 |
| CRC-2 | LN3 | 0.4993582 | 0.7382067 |
| CRC-2 | LV1 | 0.4616425 | 0.7382067 |
| CRC-2 | LV2 | 0.5741608 | 0.7382067 |
| CRC-2 | SN | 0.0091749 | 0.0825745 |
| NSCLC | P1 | 0.8600336 | 0.8838148 |
| NSCLC | P2 | 0.7366613 | 0.8838148 |
| NSCLC | P3 | 0.1747156 | 0.8838148 |
| NSCLC | C1 | 0.8219721 | 0.8838148 |
| NSCLC | C2 | 0.8838148 | 0.8838148 |
| NSCLC | C3 | 0.6516619 | 0.8838148 |
| RCC | LV1 | 0.5799825 | 0.9455936 |
| RCC | LV2 | 0.3015753 | 0.9455936 |
| RCC | LV3 | 0.9455936 | 0.9455936 |
| RCC | IVC1 | 0.9140005 | 0.9455936 |
| RCC | IVC2 | 0.7882813 | 0.9455936 |
| RCC | IVC3 | 0.5695680 | 0.9455936 |
| UBC | LN1 | 0.0109205 | 0.0174727 |
| UBC | LN2 | 0.0078581 | 0.0157162 |
| UBC | LN3 | 0.0001187 | 0.0009499 |
| UBC | U1 | 0.0032892 | 0.0131566 |
| UBC | U2 | 0.0361100 | 0.0423381 |
| UBC | P1 | 0.0055066 | 0.0146843 |
| UBC | P2 | 0.0370459 | 0.0423381 |
| UBC | P3 | 0.0553739 | 0.0553739 |

**Supplementary Table 7:**
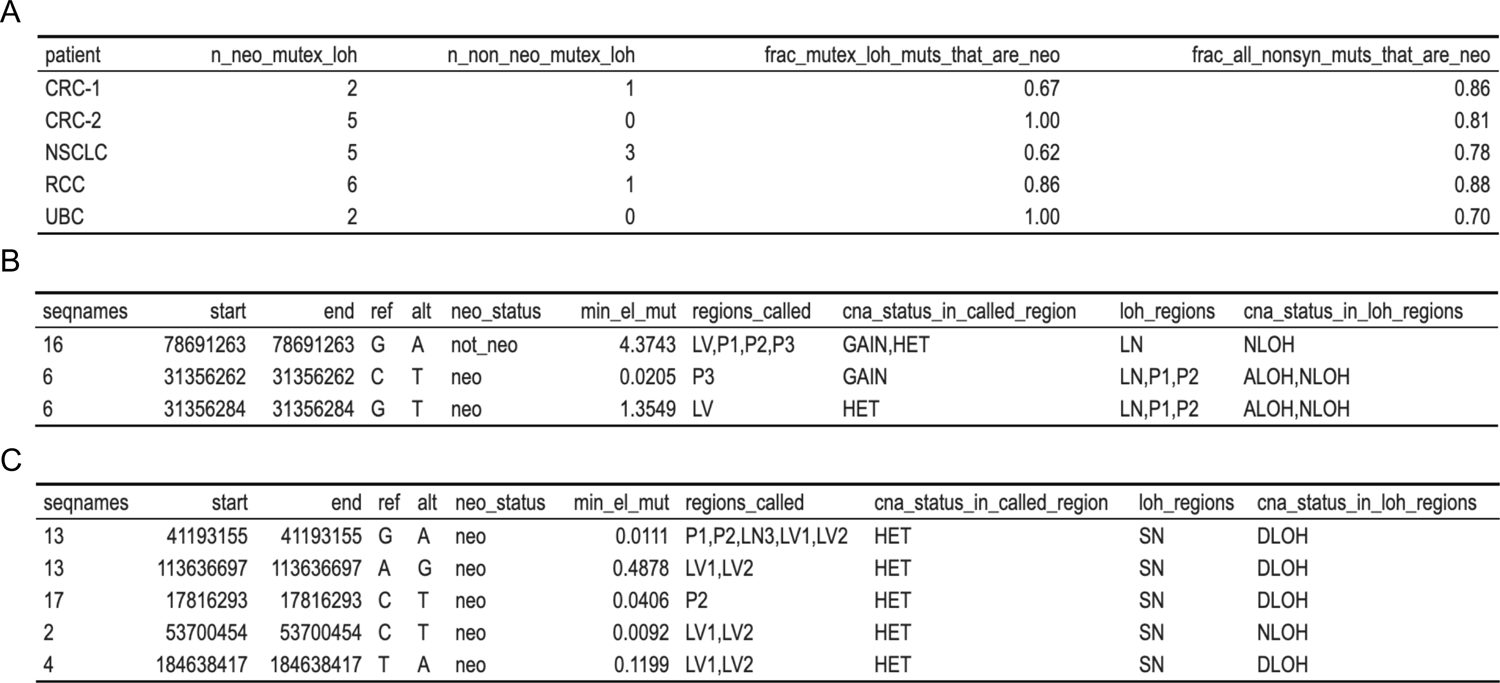
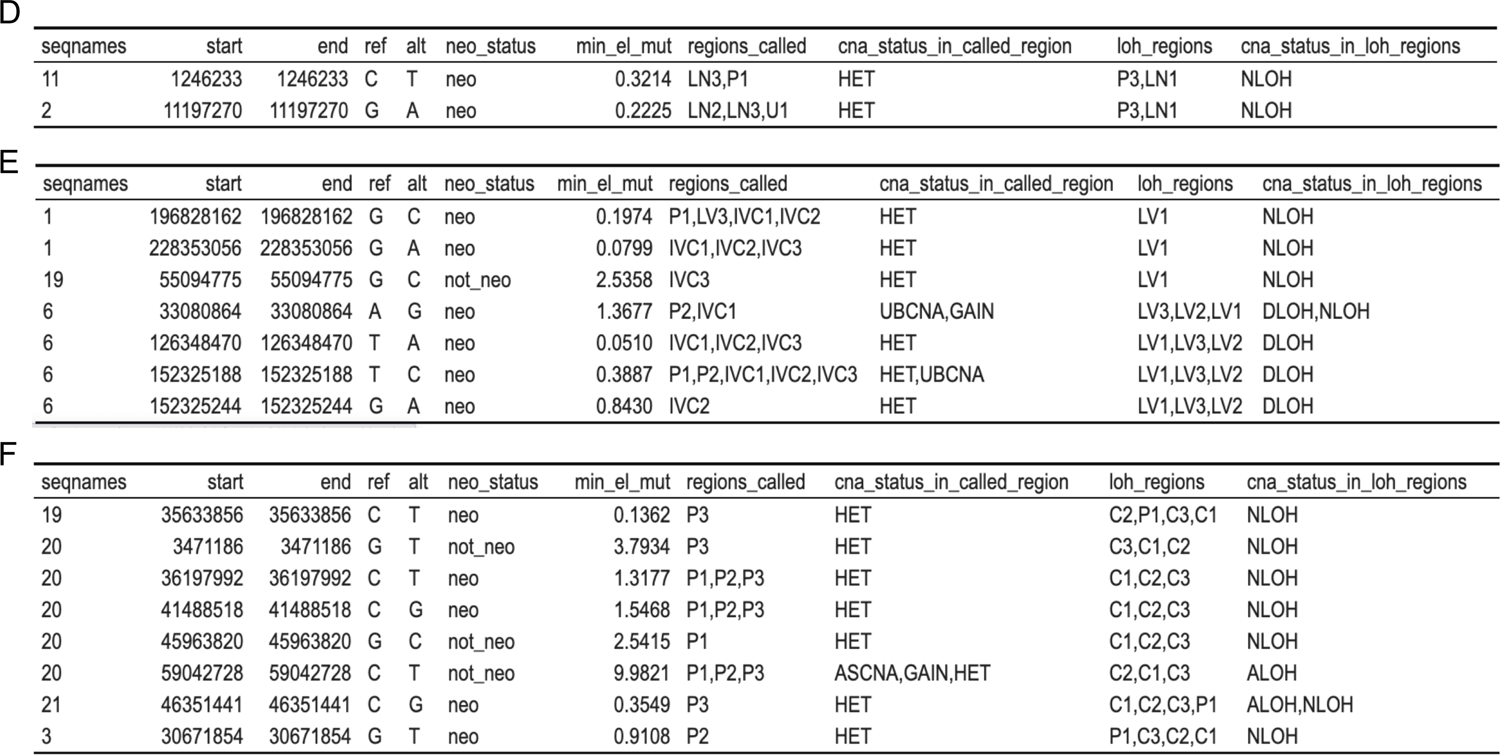
CNA based neoantigen loss in the five multi-region cases, requiring that included mutations have at least two alt-allele-supporting RNAseq reads in at least one sample. (A) Summary table showing numbers of mutations lost via genomic deletion or LOH events in each case, along with the proportion of them that are putatively neoantigenic. As a comparator, the proportion of all mutations that are putatively neoantigenic is included. (B-F) Enumeration of mutations with RNAseq support that exhibited mutual exclusivity with LOH events in individual cases - CRC-1, CRC-2, UBC, RCC, and NSCLC respectively.

Supplementary Tables 7A,B A
| patient | n_neo_mutex_loh | n_non_neo_mutex_loh | frac_mutex_loh_muts_that_are_neo | frac_all_nonsyn_muts_that_are_neo |
| --- | --- | --- | --- | --- |
| CRC-1 | 2 | 1 | 0.67 | 0.86 |
| CRC-2 | 5 | 0 | 1.00 | 0.81 |
| NSCLC | 5 | 3 | 0.62 | 0.78 |
| RCC | 6 | 1 | 0.86 | 0.88 |
| UBC | 2 | 0 | 1.00 | 0.70 |

B
| seqnames | start | end | ref | alt | neo_status | min_el_mut | regions_called | cna_status_in_called_region | loh_regions | cna_status_in_loh_regions |
| --- | --- | --- | --- | --- | --- | --- | --- | --- | --- | --- |
| 16 | 78691263 | 78691263 | G | A | not_neo | 4.3743 | LV,P1,P2,P3 | GAIN,HET | LN | NLOH |
| 6 | 31356262 | 31356262 | C | T | neo | 0.0205 | P3 | GAIN | LN,P1,P2 | ALOH,NLOH |
| 6 | 31356284 | 31356284 | G | T | neo | 1.3549 | LV | HET | LN,P1,P2 | ALOH,NLOH |

C
| seqnames | start | end | ref | alt | neo_status | min_el_mut | regions_called | cna_status_in_called_region | loh_regions | cna_status_in_loh_regions |
| --- | --- | --- | --- | --- | --- | --- | --- | --- | --- | --- |
| 13 | 41193155 | 41193155 | G | A | neo | 0.0111 | P1,P2,LN3,LV1,LV2 | HET | SN | DLOH |
| 13 | 113636697 | 113636697 | A | G | neo | 0.4878 | LV1,LV2 | HET | SN | DLOH |
| 17 | 17816293 | 17816293 | C | T | neo | 0.0406 | P2 | HET | SN | DLOH |
| 2 | 53700454 | 53700454 | C | T | neo | 0.0092 | LV1,LV2 | HET | SN | NLOH |
| 4 | 184638417 | 184638417 | T | A | neo | 0.1199 | LV1,LV2 | HET | SN | DLOH |

Supplementary Tables 7D,E,F D
| seqnames | start | end | ref | alt | neo_status | min_el_mut | regions_called | cna_status_in_called_region | loh_regions | cna_status_in_loh_regions |
| --- | --- | --- | --- | --- | --- | --- | --- | --- | --- | --- |
| 11 | 1246233 | 1246233 | C | T | neo | 0.3214 | LN3,P1 | HET | P3,LN1 | NLOH |
| 2 | 11197270 | 11197270 | G | A | neo | 0.2225 | LN2,LN3,U1 | HET | P3,LN1 | NLOH |

E
| seqnames | start | end | ref | alt | neo_status | min_el_mut | regions_called | cna_status_in_called_region | loh_regions | cna_status_in_loh_regions |
| --- | --- | --- | --- | --- | --- | --- | --- | --- | --- | --- |
| 1 | 196828162 | 196828162 | G | C | neo | 0.1974 | P1,LV3,IVC1,IVC2 | HET | LV1 | NLOH |
| 1 | 228353056 | 228353056 | G | A | neo | 0.0799 | IVC1,IVC2,IVC3 | HET | LV1 | NLOH |
| 19 | 55094775 | 55094775 | G | C | not_neo | 2.5358 | IVC3 | HET | LV1 | NLOH |
| 6 | 33080864 | 33080864 | A | G | neo | 1.3677 | P2,IVC1 | UBCNA,GAIN | LV3,LV2,LV1 | DLOH,NLOH |
| 6 | 126348470 | 126348470 | T | A | neo | 0.0510 | IVC1,IVC2,IVC3 | HET | LV1,LV3,LV2 | DLOH |
| 6 | 152325188 | 152325188 | T | C | neo | 0.3887 | P1,P2,IVC1,IVC2,IVC3 | HET,UBCNA | LV1,LV3,LV2 | DLOH |
| 6 | 152325244 | 152325244 | G | A | neo | 0.8430 | IVC2 | HET | LV1,LV3,LV2 | DLOH |

F
| seqnames | start | end | ref | alt | neo_status | min_el_mut | regions_called | cna_status_in_called_region | loh_regions | cna_status_in_loh_regions |
| --- | --- | --- | --- | --- | --- | --- | --- | --- | --- | --- |
| 19 | 35633856 | 35633856 | C | T | neo | 0.1362 | P3 | HET | C2,P1,C3,C1 | NLOH |
| 20 | 3471186 | 3471186 | G | T | not_neo | 3.7934 | P3 | HET | C3,C1,C2 | NLOH |
| 20 | 36197992 | 36197992 | C | T | neo | 1.3177 | P1,P2,P3 | HET | C1,C2,C3 | NLOH |
| 20 | 41488518 | 41488518 | C | G | neo | 1.5468 | P1,P2,P3 | HET | C1,C2,C3 | NLOH |
| 20 | 45963820 | 45963820 | G | C | not_neo | 2.5415 | P1 | HET | C1,C2,C3 | NLOH |
| 20 | 59042728 | 59042728 | C | T | not_neo | 9.9821 | P1,P2,P3 | ASCNA,GAIN,HET | C2,C1,C3 | ALOH |
| 21 | 46351441 | 46351441 | C | G | neo | 0.3549 | P3 | HET | C1,C2,C3,P1 | ALOH,NLOH |
| 3 | 30671854 | 30671854 | G | T | neo | 0.9108 | P2 | HET | P1,C3,C2,C1 | NLOH |

